# Planar cell polarity in the larval epidermis of *Drosophila* and the role of microtubules

**DOI:** 10.1101/2020.02.21.959213

**Authors:** Stefano Pietra, KangBo Ng, Peter A. Lawrence, José Casal

**Affiliations:** Department of Zoology, University of Cambridge, Downing Street, Cambridge CB2 3EJ, United Kingdom; The Francis Crick Institute, 1 Midland Road, London NW1 1AT and Institute for the Physics of Living Systems, University College London, London, United Kingdom

**Keywords:** Planar Cell Polarity, *Drosophila*, larval epidermis, microtubules, protocadherins, *dachsous*, *fat*, *ovo*, *dachs*

## Abstract

We investigate the mechanisms of planar cell polarity (PCP) in the *Drosophila* larva. The epidermis displays an intricate pattern of polarity and is excellent for the study of one system of PCP, the Dachsous/Fat system; partly because the Starry Night/Frizzled system plays no discernable role in the larva. Measurements of the amount of Dachsous reveal a peak near the rear of the anterior compartment. Localisation of Dachs and orientation of ectopic denticles reveal the polarity of every cell in the segment. We discuss how well these findings evidence our gradient model of Dachsous activity. Several groups have proposed that Dachsous and Fat fix the direction of PCP via oriented microtubules that transport PCP proteins to one side of the cell. We test this proposition in the larval cells and find that most microtubules grow perpendicularly to the axis of PCP. We find no meaningful bias in the polarity of those microtubules aligned close to that axis. We also reexamine published data from the pupal abdomen and fail to find evidence supporting the hypothesis that microtubular orientation draws the arrow of PCP.

## INTRODUCTION

As cells construct embryos and organs they need access to vectorial information that informs them, for example, which way to migrate, divide, extend axons and orient protrusions such as hairs. This kind of polarity is known as planar cell polarity (PCP). In *Drosophila* there are (at least) two conserved genetic systems that generate PCP. Both systems rely on the formation of intercellular bridges made by transmembrane proteins containing cadherin repeats, these interact via their extracellular domains. The Dachsous/Fat (Ds/Ft) system depends on heterodimers of the protocadherins Ds and Ft while the Starry Night/Frizzled system relies on homodimers of Starry Night (reviewed in [**1-6**]). Most developmental models can be tricky to study because both PCP systems operate at once and both have separate but confounding inputs into the orientation of bristles, etc. However, here we investigate the later stage larvae in which PCP depends entirely on the Ds/Ft system [**7-9**] whose mechanism is quite well understood. Ds molecules in one cell bind to Ft molecules in a neighbour cell to make intercellular bridges. Experiments argue that, using the disposition and orientation of Ds-Ft bridges, each cell compares the Ds activity of those two of its neighbours that lie in the relevant axis and points its denticles towards the neighbour with the ***higher*** Ds activity. Ds activity is thus an important component of the model: the activity of Ds in a cell defines its ability to bind to Ft in its neighbouring cell, that activity depending on at least three factors; the levels of Ds expression, the levels of Ft expression and the activity of Four-jointed (Fj). Fj is a Golgi-resident kinase that phosphorylates both Ds and Ft, reducing the activity of the former while increasing the activity of the latter [**10-12**].

The system has an additional property: because of the interdependence of membrane bound Ds and Ft in neighbouring cells, the polarity of one cell can affect the polarity of its neighbours and that polarity can be propagated to the next neighbour [**7, 13, 14**]. Thus, in these several ways the landscape of Ds activity in a field of cells is translated into the individual polarities of the cells (see [**5**] for further explanation). More recently, we have, via experiments and observations, developed a model that explains the quite complex pattern of denticle polarities in the larval abdominal segment [**15**].

### A model: the ventral epidermis of the *Drosophila* larva

Each segment of the larva is divided by cell lineage into an anterior (A) and a posterior (P) compartment. In the adult abdomen, the A and P compartments are thought to be approximately coextensive with opposing gradients of Ds activity [**16**] and if such gradients were present in the larva then they could explain most of the denticle polarities. However, in the larva, in addition to the normal denticulated cells, there are three interspersed rows of muscle attachment cells [**15, 17, 18**] and our experiments suggest that two of these three rows have exceptionally low Ds activity which can affect the polarity of neighbouring cells (**figure 1**, [**15, 17**]). At this point we are not clear how much the final pattern is determined by pervasive gradients of Ds activity or how much by these local effects of the muscle attachment cells plus propagation.

**Figure 1.**
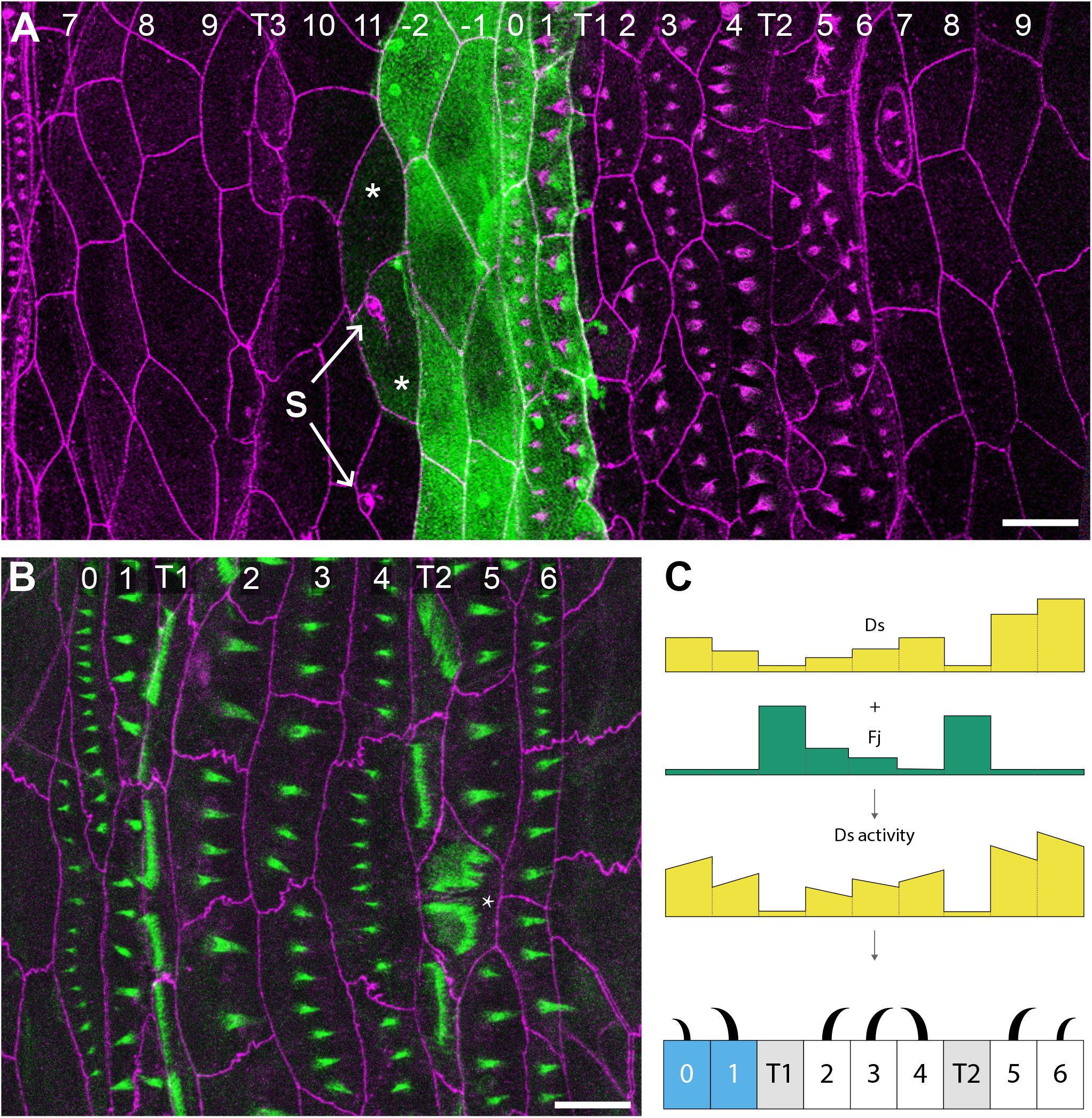
Larval ventral abdomen and Ds activity landscape. (**A**) Overview of segments with cells expressing GFP under the control of the *engrailed* promoter, a marker of the P compartment [**51, 58**]. GFP labels four rows of cells, between the most posterior row of the A compartment (identified by sensory cells, S) and the most anterior row of the following segment (tendon cells T1, see [**18**]). This driver occasionally also weakly labels a few cells at the rear of the A compartment (asterisks), but we have found that these cells do not express other P markers such as *hedgehog* (data not shown). Cell outlines and denticles are labelled in magenta (DE-cad::tomato). Arrows point to sensory cells (s) that we used as positional markers. (**B**) Ventral denticulate area of a mid second stage larva. Predenticles (rows 0 to 6) and tendon cells (rows T1 and T2) are marked in green (UTRN::GFP, labelling actin), and cell boundaries in magenta (DE-cad::tomato). The rows are not completely regular; here, one T2 cell contacts two row 6 cells at the posterior (asterisk) — typically, T2 only contacts row 5 cells. (**C**) A partially documented model of the landscape of Ds and Fj and therefore of PCP in the wild type [**15, 17**]. In this model, a presumed low level of *ds* expression together with a documented high level of Fj reduces Ds activity in T1 and T2. The sloped line in each cell indicates different amounts of Ds activity at its anterior and posterior limits, the direction of the slope correlating with the cell’s polarity. Denticle polarity is shown below and is a readout of the presumed landscape of Ds activity: each cell points its denticles towards the neighbour with the higher Ds activity. Two rows of the P compartment are highlighted in blue, tendon cells are shaded in grey. Anterior is to the left in all figures. Scale bars: 20μm.

One outstanding difficulty in applying present models to the whole segment is that more than half the cells do not make denticles and their polarities are not known. In this paper we have solved that difficulty by measuring the molecular polarities of these uncharted cells in two complementary and different ways and this allows us to extend model-building to the entire segment. With the same purpose we have also measured the amount of Ds expression in each intercellular junction across the entire segment.

Depending on the pattern of Ds activity, individual cells will acquire different numbers of Ds-Ft and Ft-Ds heterodimers at opposite cell faces. Generally this difference will explain the polarity of the whole cell, however, sometimes and depending on the disposition of neighbouring cells, two regions of a single cell can have opposing polarities [**17**]. To explain this phenomenon it has been argued that polarity of individual cells or parts of cells would depend on local “conduits” that run between opposing cell faces to mediate their comparison. In this paper we reinvestigate these multipolar cells in an experimental situation.

There is some evidence that suggests that these conduits acting within the Ds/Ft system could be microtubules and might polarise the cell by orienting the intracellular transport of molecules and vesicles [**19, 20**]. Indeed Harumoto et al reported that, in one particular region of the pupal wing, the majority of microtubules are aligned near-parallel with the axis and direction of PCP (the direction of PCP is defined by the orientation of hairs) and, when growing, they show a small but statistically significant “bias” in polarity [**20**]. By bias we mean a net difference between the number of microtubules growing within a particular angle interval and the number of microtubules growing 180 degrees away; for instance we might see more microtubules growing distally, ie in the same direction as the hairs, than in the opposite direction. Harumoto et al therefore proposed that, in general, the Ds/Ft system controls the orientation of microtubules that would subsequently polarise cells by serving as oriented conduits in the polarised transport of PCP components [**20**]. Tests of this hypothesis in the adult abdomen have given mixed results [**21-23**]. Results from both wing and the abdomen are conflicting; regions of both appear to be polarised independently of the microtubules [**23**]. In the hope of clarifying this confusing situation we now report our studies of microtubule orientation *in vivo* in the larva. The larva has some advantages over imaginal discs or the adult abdomen: individually identifiable cells have a defined polarity and larval cells are much larger than the adult cells allowing more precision in plotting of the orientation of the microtubules. Several analyses of our own results on the larval abdomen and of raw data kindly provided by Axelrod from the pupal abdomen [**22, 23**] do not support the hypothesis that PCP is oriented by microtubules.

In this paper we add to our knowledge of PCP in the larval segment; our two most important findings are to define cell polarity in all the cells of the entire segment and to provide data arguing strongly that orientation of the microtubules does not correlate with the axis of denticle polarity.

## MATERIALS AND METHODS

### Mutations and Transgenes

Flies were reared at 25°C on standard food. The FlyBase [**24**] entries for the mutant alleles and transgenes used in this work are the following: *ds*: *ds*^*UA071*^; *en*.*Gal4*: *Scer\GAL4*^*en-e16E*^; *sr*.*Gal4*: *sr*^*md710*^; *UAS*.*act*::*GFP*: *Dmel\Act5C*^*UAS*.*GFP*^; *UAS*.*DsRed*: *Disc\RFP*^*UAS*.*cKa*^; *UAS*.*EB1*::*EGFP*: Eb1^UAS.GFP^; *UAS*.*ectoDs*: *ds*^*ecto*.*UAS*^; *UAS*.*LifeAct*::*mCherry*: *Scer\ABP140*^*UAS*.*mCherry*^; *UAS*.*RedStinger: Disc\RFP*^*DsRedT4*.*UAS*.*Tag:NLS(tra)*^; *UAS*.*ovo*: *ovo*^*svb*.*Scer\UAS*^; *act>stop>d*::*EGFP*: *d*^*FRT*.*Act5C*.*EGFP*^; *DE-cad*::*tomato*: *shg*^*KI*.*T:Disc\RFP-tdTomato*^; ds::EGFP: *Avic\GFP*^*ds-EGFP*^; *hs*.*FLP*: *Scer\FLP1*^*hs*.*PS*^; *sqh*.*UTRN*::*GFP*: *Hsap\UTRN*^*Scer\UAS*.*P\T*.*T:Avic\GFP-EGFP*^; *tub>stop>Gal4*: *Scer\GAL4*^*FRT*.*Rnor\Cd2*.*αTub84B*^.

### Experimental Genotypes

(**figure 1A)** *y w hs*.*FLP/ w; DE-cad::tomato/ en*.*Gal4 UAS*.*act::GFP*.

**(figure 1B)** *w; DE-cad::tomato sqh*.*UTRN::GFP*.

**(figure 2**, and **table 1)** *w; ds*^*UA071*^ *DE-cad::tomato sqh*.*UTRN::GFP/ DE-cad::tomato sqh*.*UTRN::GFP; sr*.*Gal4/ UAS*.*ectoDs*.

**Table 1.**
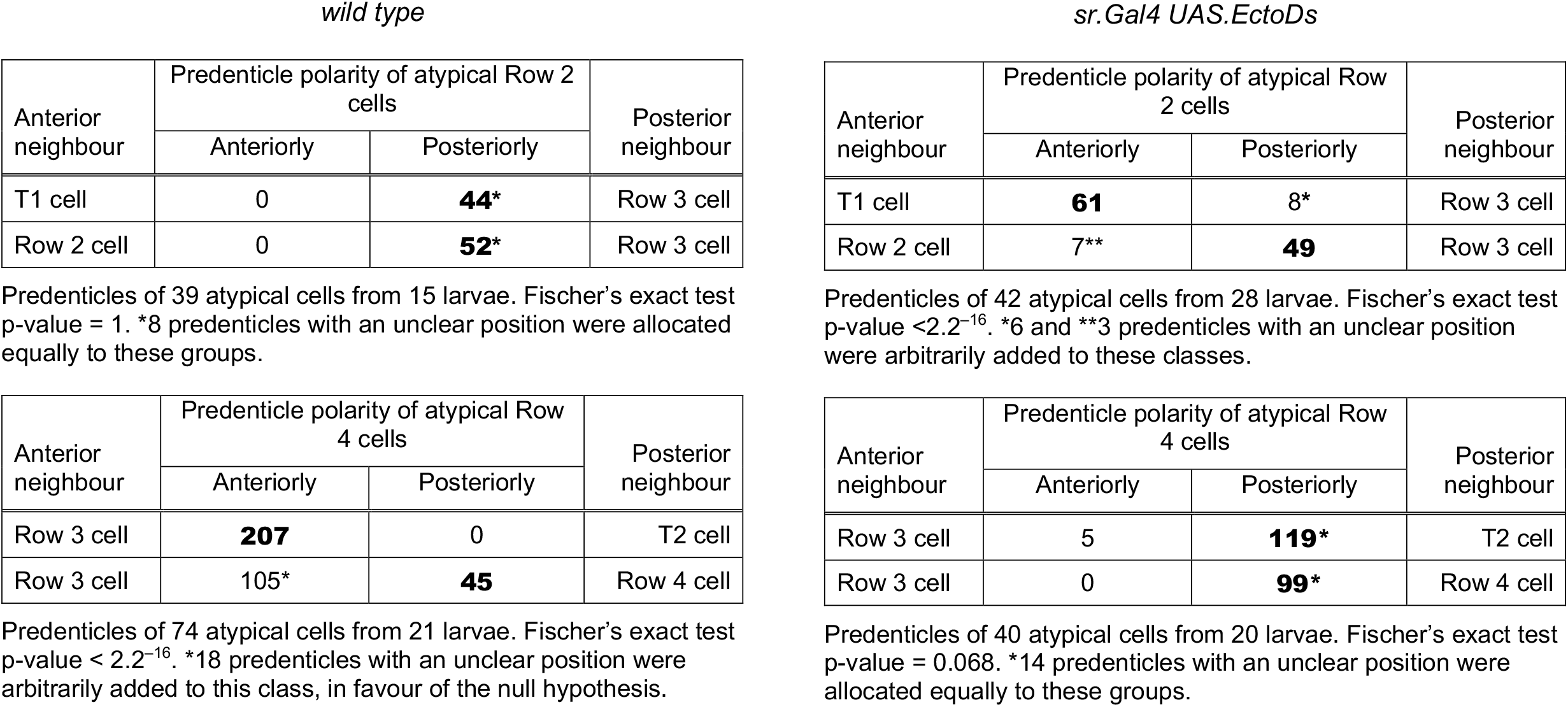
Atypical cells: quantitation of predenticle polarities in relation to neighbouring cells, showing the effect of over expressing *ds* in the Tendon cells.

| Anterior neighbour | Predenticle polarity of atypical Row 2 cells |  | Posterior neighbour |
| --- | --- | --- | --- |
|  | Anteriorly | Posteriorly |  |
| T1 cell | 0 | **44\*** | Row 3 cell |
| Row 2 cell | 0 | **52\*** | Row 3 cell |

| Anterior neighbour | Predenticle polarity of atypical Row 4 cells |  | Posterior neighbour |
| --- | --- | --- | --- |
|  | Anteriorly | Posteriorly |  |
| Row 3 cell | **207** | 0 | T2 cell |
| Row 3 cell | 105* | **45** | Row 4 cell |

| Anterior neighbour | Predenticle polarity of atypical Row 2 cells |  | Posterior neighbour |
| --- | --- | --- | --- |
|  | Anteriorly | Posteriorly |  |
| T1 cell | **61** | 8* | Row 3 cell |
| Row 2 cell | 7** | **49** | Row 3 cell |

| Anterior neighbour | Predenticle polarity of atypical Row 4 cells |  | Posterior neighbour |
| --- | --- | --- | --- |
|  | Anteriorly | Posteriorly |  |
| Row 3 cell | 5 | **119\*** | T2 cell |
| Row 3 cell | 0 | **99\*** | Row 4 cell |

Predenticles of 40 atypical cells from 20 larvae. Fischer's exact test p-value = 0.068. \*14 predenticles with an unclear position were allocated equally to these groups.

**Figure 2.**
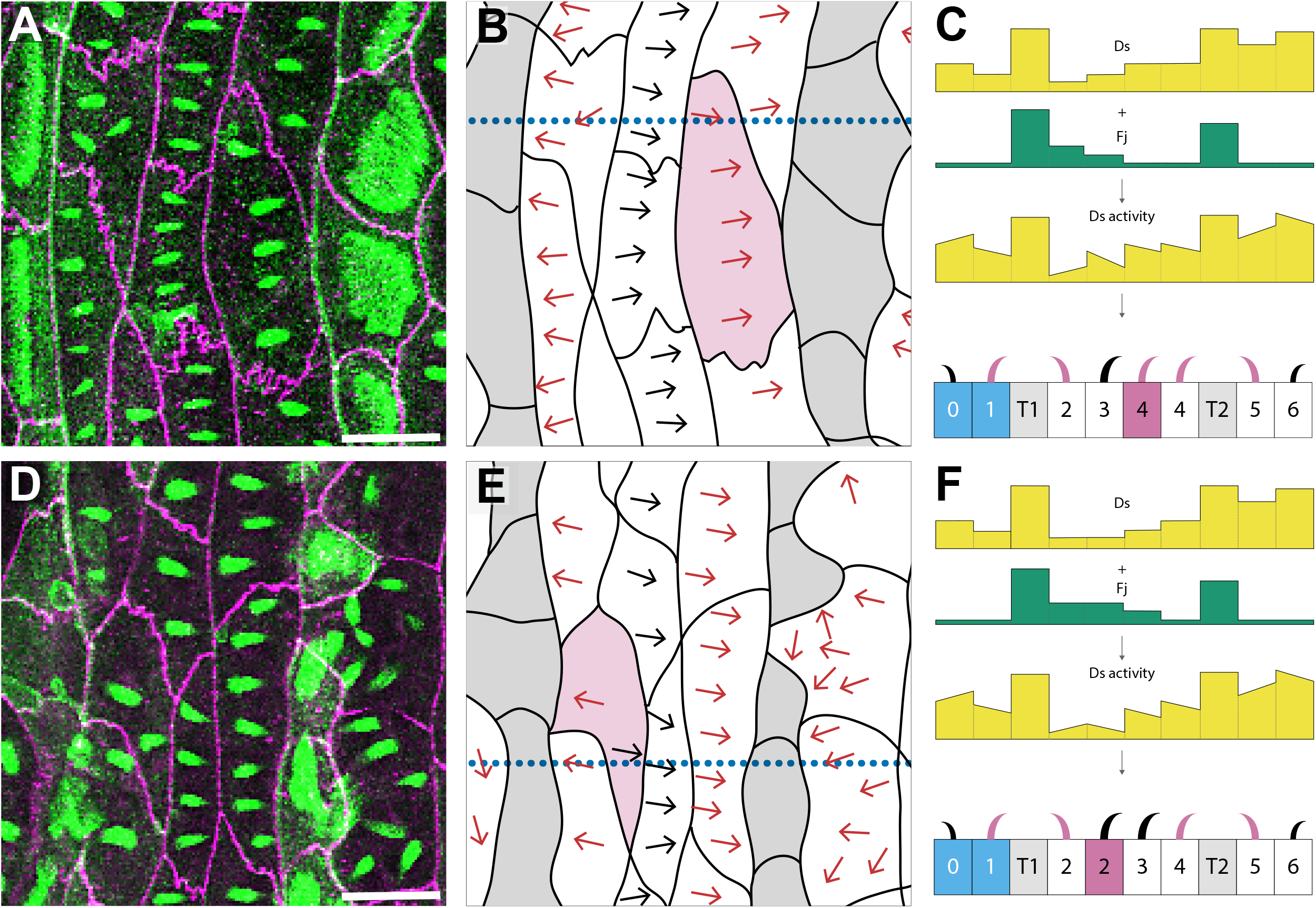
PCP and atypical cells in polarity modified larvae. Denticulate areas of polarity modified larvae: (**A-C**) an atypical cell in row 4 (having two posterior neighbours with different Ds activity), and (**D-F**) an atypical cell in row 2 (having two anterior neighbours with different Ds activity). Predenticles and denticles in rows 1, 2 and 4, 5 with polarity opposite from wildtype are highlighted in magenta. (**A,D**) Images of predenticles, tendon cells, and cell boundaries labelled as in **figure 1B**. (**B,E**) Schemes of cell outlines and predenticle orientation. (**C,F**) Models of polarity modified larvae, Ds activity landscape and denticle polarity in cross sections taken at the dotted blue lines in **B,E**. Blue shading indicates P compartment cells, grey denotes tendon cells, magenta marks the atypical cell. Note that, contrary to wildtype [**17**], in polarity modified larvae row 4 atypical cells are monopolar (**A,B**), while row 2 atypical cells are multipolar (**D,E**). For quantitation of predenticle polarity in row 4 and row 2 atypical cells of wild type and polarity modified larvae, see **Table 1**. Scale bars: 20μm.

**(figures 3, 4)** *w; ds::EGFP FRT40A*.

**Figure 3.**
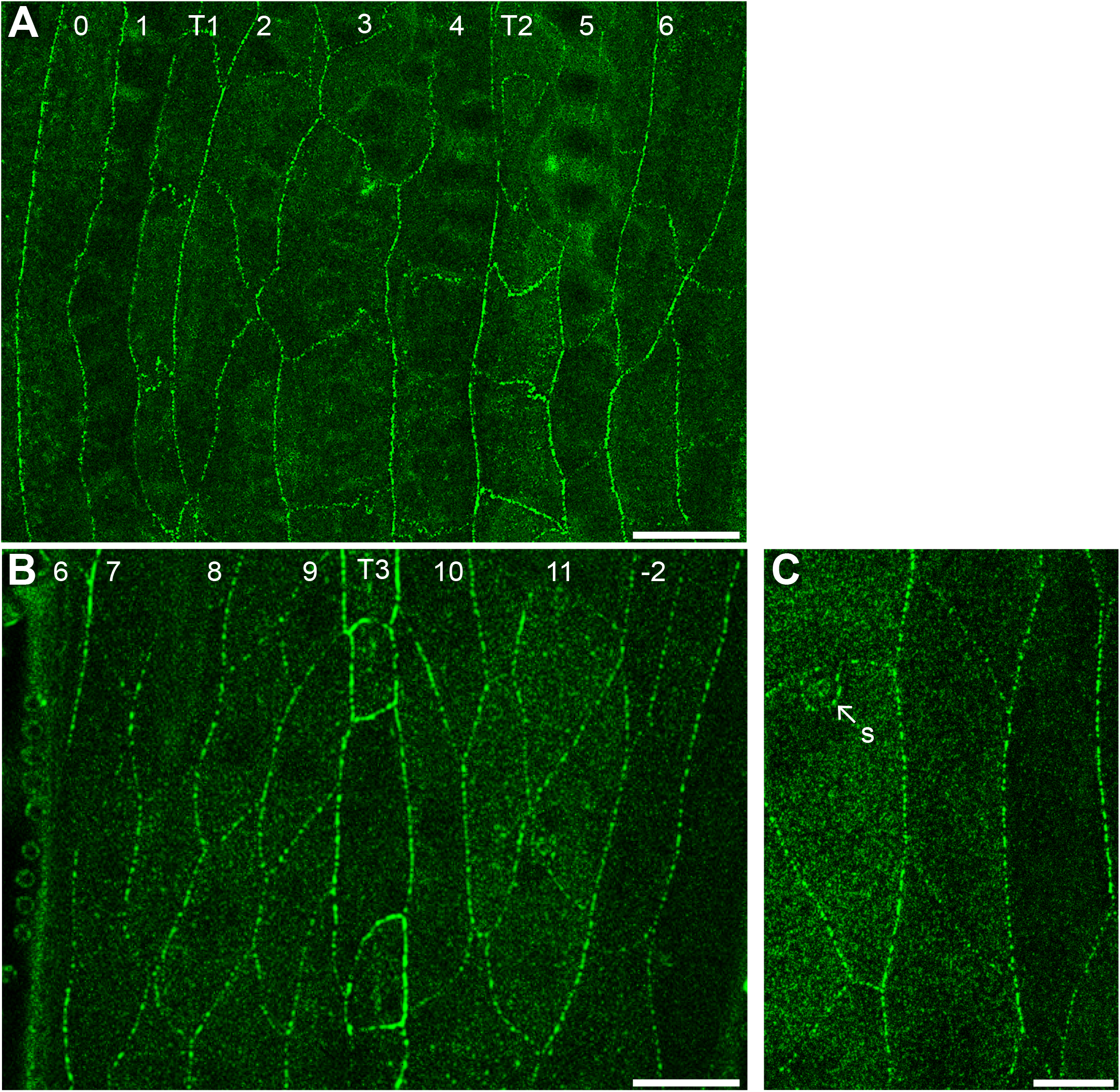
Ds localisation in the larval ventral abdomen. Larvae expressing *ds*::EGFP from the tagged endogenous *ds* locus [**14**] show a ubiquitous punctate pattern of fluorescence that concentrates on plasma membranes. (**A**) Denticulate and (**B**) undenticulate areas of early second stage larvae; the cell rows exhibit no obvious differences in *ds* expression or distribution, with the exception of the strong signal around T3 tendon cells. (**C**) Detail of Ds localisation in puncta at the cell membrane. 0 to 6, denticle cell rows. 7 to −2, undenticulate cell rows. S, sensory cell. T1, T2, T3, tendon cell rows. Scale bars: 20μm (**A,B**), 10μm (**C**).

**Figure 4.**
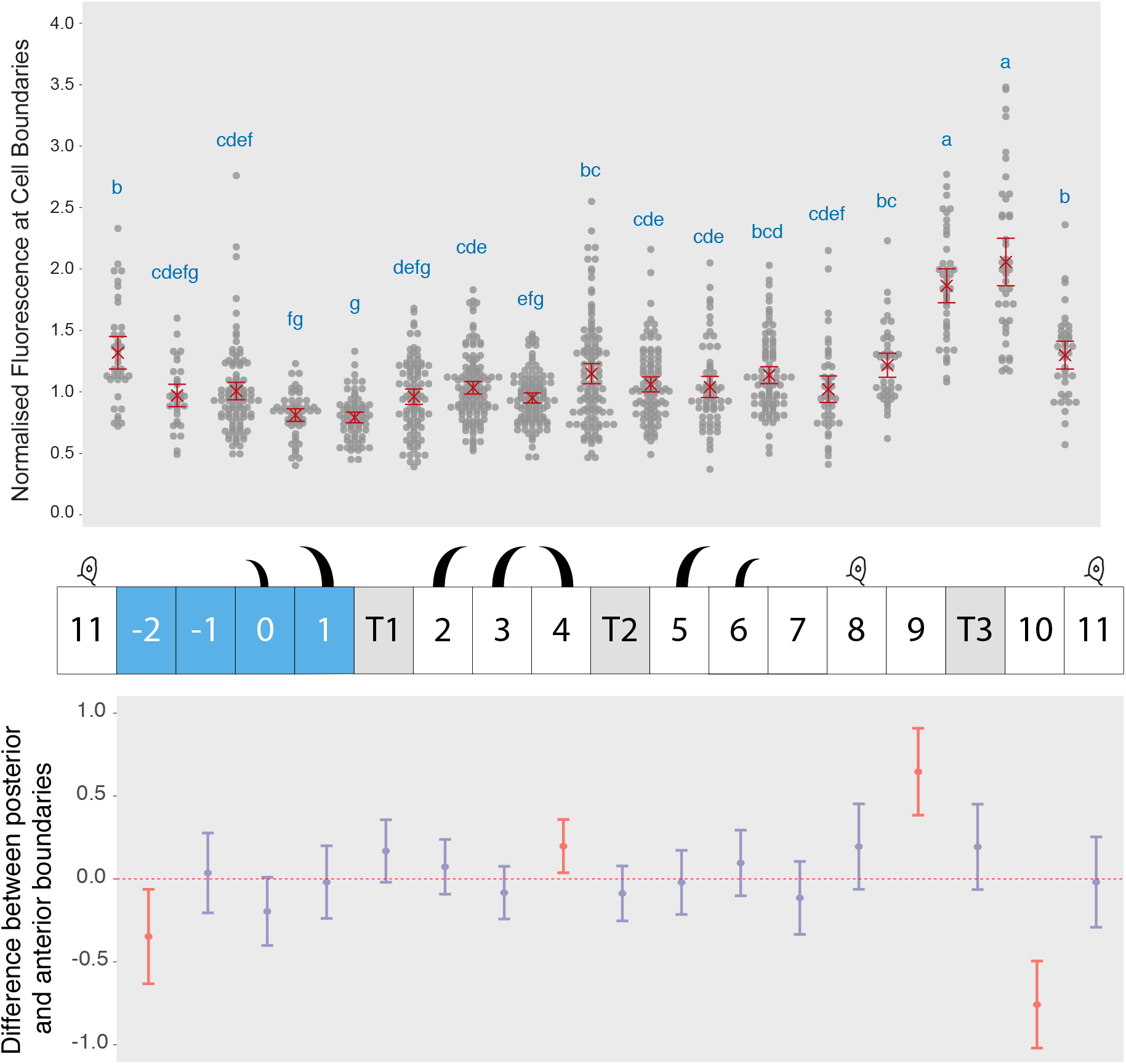
Quantitation of Ds levels at cellular interfaces across the segment. (**Top**) Dot plot of normalised fluorescence intensity maxima corresponding to amounts of Ds at boundaries between cell rows of the larval ventral abdomen. Data are pooled from 12 (denticulate area) and 5 (undenticulate area) images of different larvae. Mean value and 95% confidence interval for each interface are indicated in red. Letters arise from Tukey’s multiple comparison test between all interfaces; in the Tukey’s test, comparisons between pairs belonging to a group with the same letter show a p value equal to or greater than 0.05. Groups can be assigned more than one letter, reflecting “overlap” between different groups. The graph shows no evidence for a segment-wide gradient of Ds accumulation at the cell membranes, however the 9/T3 and T3/10 boundaries are significantly different from all others, indicating a clear peak anterior to the A/P boundary. (**Middle**) Diagram of denticle polarity, as in **figure 1C**. Sensory cells identify rows 8 and 11. (**Bottom**) Comparisons between Ds amounts at posterior and anterior interfaces of each cell row. Differences in mean normalised fluorescence at the opposite sides of a cell are calculated with 95% confidence interval by Tukey’s test. Red indicates a significant difference. Note the significant and opposite differences in cell rows 9 and 10, highlighting the presence of a fluorescence peak around T3.

**(figures 5A,B, 6)** *y w hs*.*FLP/ w; en*.*Gal4 UAS*.*DsRed/ +; act>stop>d::EGFP/ +*.

**Figure 5.**
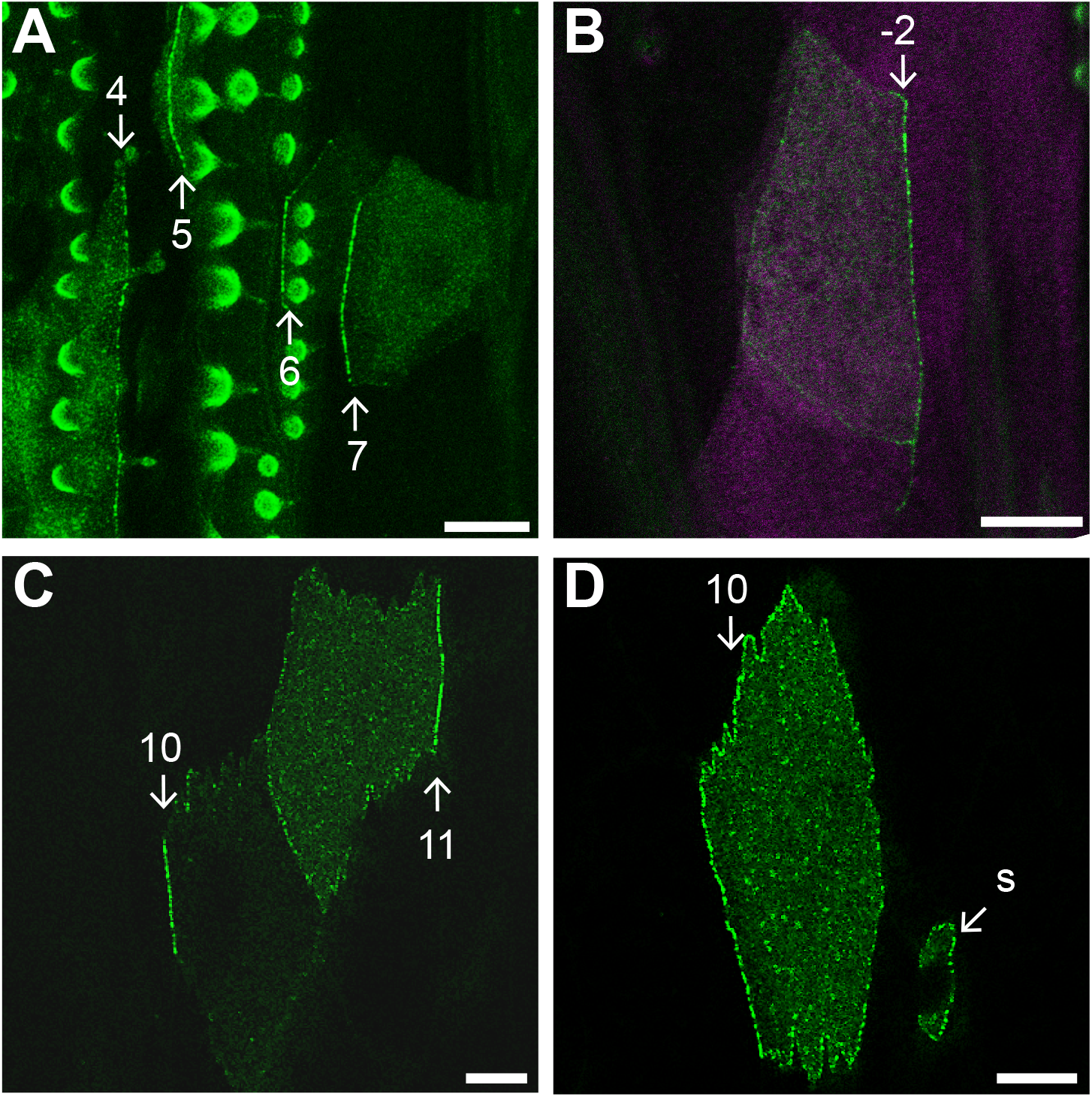
D polarity at the plasma membrane in small clones. (**A**) Several cells of the A compartment expressing *d::EGFP*: in row 4, where denticles point anteriorly, D is mostly on the posterior membrane; in rows 5, 6 and 7, with posterior-pointing polarity, D accumulates instead at the anterior face of the cells. Round or comma-like structures are due to autofluorescence from overlying denticles. (**B**) A posterior cell (row −2) accumulates D at its rear, arguing for anterior-pointing polarity. P compartment is labelled in magenta by *en*.*Gal4 UAS*.*DsRed*. (**C**) Cells of rows 10 and 11, where D localises on the anterior and posterior sides of the plasma membrane, respectively (see **figure S2** for cell outlines). (**D**) Row 10 cell with more D on the anterior side of the cell membrane, suggesting its polarity points backwards. The sensory cell process associated with row 11 also expresses *d::EGFP*, and as with other cells from row 11 has most D at the posterior side. S, sensory cell. Scale bars: 10μm.

**Figure 6.**
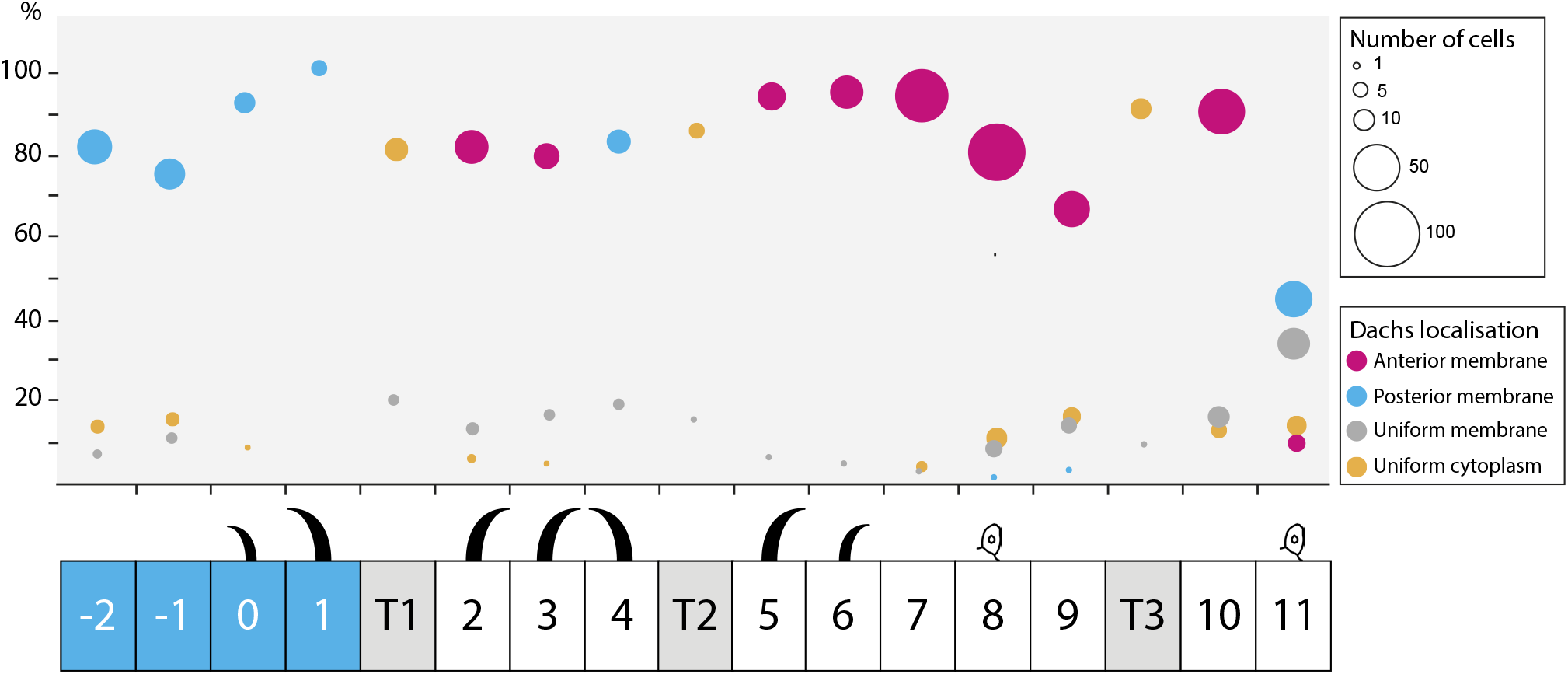
The localisation of D cell by cell. D localisation in all the cell rows, derived from the analysis of small clones expressing *d::EGFP*. Cells where D accumulates on just the anterior side of the plasma membrane contribute to red circles (Anterior membrane), cells where D is only on the posterior side to blue circles (Posterior membrane), cells where D is enriched at the plasma membrane but in an unpolarised manner to grey circles (Uniform membrane), and cells where D is homogeneously distributed in the cytoplasm to orange circles (Uniform cytoplasm). The position of each circle denotes the cell row and percentage of cells with the indicated D localisation in that row; circle area is proportional to the number of cells represented. Since D is thought to accumulate on the side of a cell facing the neighbour with the least Ds, the pattern of D polarity in the undenticulate region suggests that there is a peak of Ds activity in row 10 (see **figure 9** for full model). n = 594 cells from a total of 44 larvae.

**(figures 5C,D, S2, 6)** *y w hs*.*FLP/ w; DE-cad::tomato; act>stop>d::EGFP/ +*.

**(figures 7, S3)** *y w hs*.*FLP/ w; tub>stop>Gal4/ DE-cad::tomato; UAS*.*ovo/ UAS*.*EB1::EGFP*.

**Figure 7.**
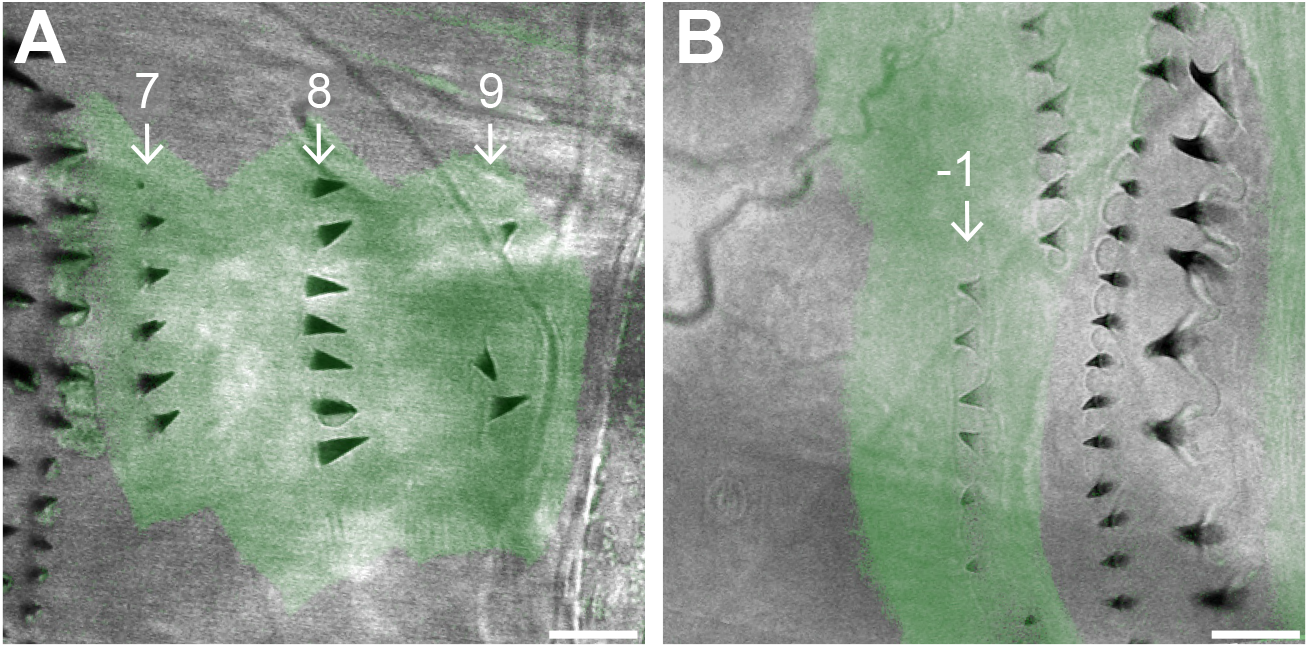
*ovo*-overexpressing clones in normally undenticulate areas of the epidermis. (**A**) Clone in the A compartment (cell rows 7, 8, and 9), marked with EGFP and producing ectopic denticles that point backwards. (**B**) Clone in the P compartment (cell row −1), ectopic denticles pointing forwards. Note that denticles are produced somewhat sporadically and that denticle numbers vary per cell. Scale bars: 10μm.

**(figures 8, S4, S6**, and **movies 1, 2)** *y w hs*.*FLP/ w; tub>stop>Gal4/ DE-cad::tomato; UAS*.*EB1::EGFP/ UAS*.*LifeAct::mCherry*.

**Figure 8.**
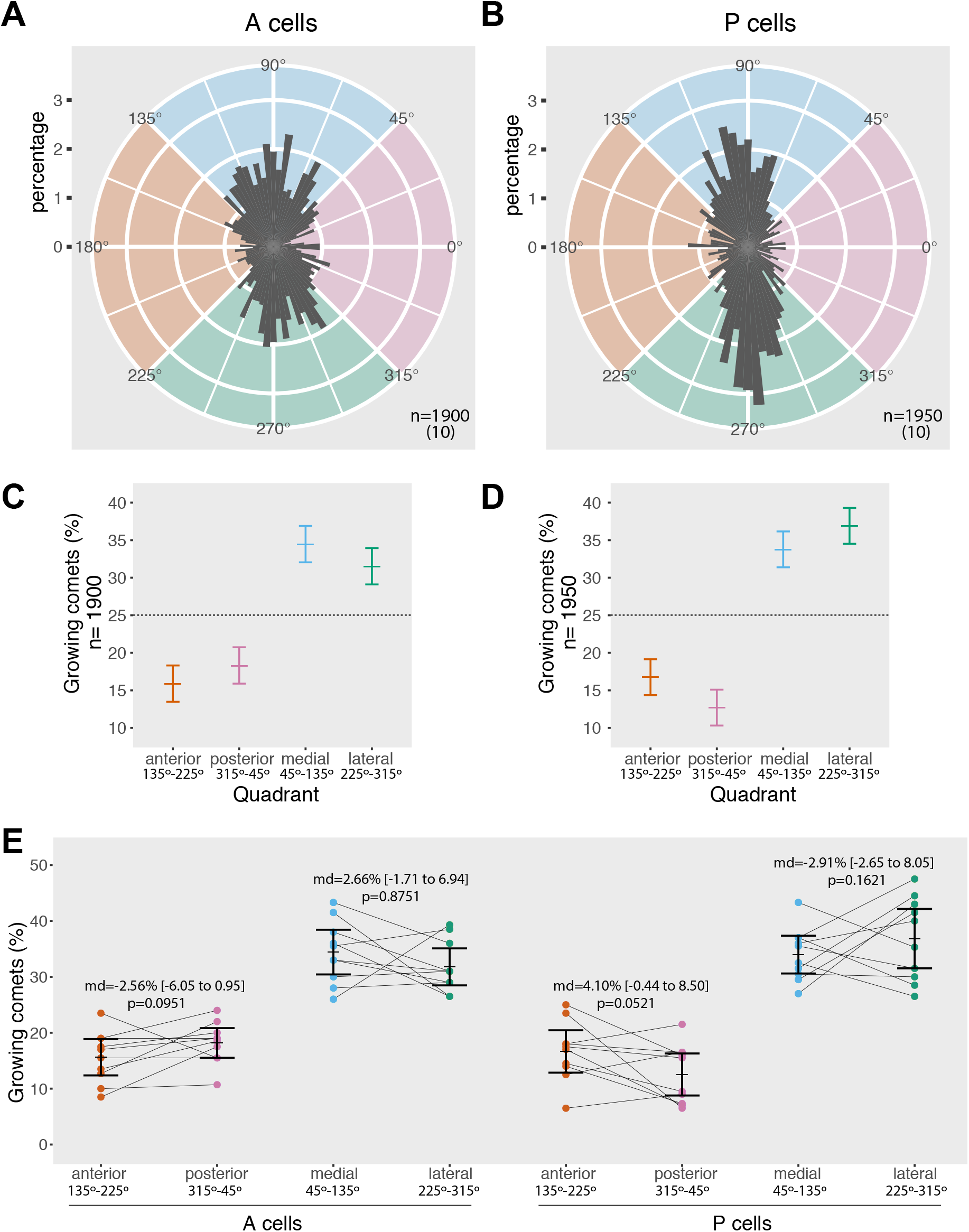
Analysis of microtubule polarity in larval epidermal cells. (**A,B**) Rose diagrams showing the distribution of growing microtubule direction in cells of the (**A**) anterior and (**B**) posterior compartment. EB1 comets are grouped in bins of 4 degrees, the length of each bin indicating the percentage of comets with a specific orientation. Comets pointing to the left (135-225°, orange quadrant) grow anteriorly, comets pointing to the right (315-45°, pink) posteriorly, up (45-135°, blue) are medial, and down (225-315°, green) are lateral; n is the total number of comets tracked, from the number of cells/larvae indicated in parenthesis. (**C,D**) Frequency of microtubules with either anterior, posterior, medial or lateral orientation in (**C**) A cells and (**D**) P cells. Comets are sorted into four sectors of 90 degrees centred on the anteroposterior and mediolateral axes. The 95% confidence interval for all comets in each quadrant is calculated according to Sison and Glaz [**59**]. (**E**) Dot plot comparing the orientation of microtubules within each cell of the A and P compartment. For every cell, the fraction of comets falling into the anterior quadrant is plotted next to the fraction in the posterior quadrant, medial next to lateral. Lines connecting the twin values from the same cell emphasise the high variability between individuals. Mean percentage and 95% confidence interval of the mean for each set of cells are shown. Overlying numbers display the exiguous difference between means (md) of the anterior versus posterior and medial versus lateral quadrants, with 95% confidence interval estimated by recalculating the difference of the means after resampling the data 10,000 times and finding the 0.025 and 0.975 quantiles of the resulting distribution of values; P-values were obtained as the frequency of resampled differences of the means that were greater than the observed.

**(figure S1)** *w; ds::EGFP FRT40A/ +; UAS*.*ectoDs/ sr*.*Gal4 UAS*.*RedStinger*.

### Live Imaging of Larvae

To induce clones expressing *d::EGFP, ovo*, or *EB1::EGFP*, 2-4 h AEL embryos were heat shocked on agar plates with fresh yeast paste at 33°C for 30 min in a water bath. Larvae were grown at 25°C for 47-52 hr and moved to fresh standard food for 2-4 h (tagged Ds, D, and EB1) or 10-15 hr (predenticles) before imaging. Second stage larvae were washed in water and then immobilised between a glass slide and coverslip by exploiting the surface tension of a drop of Voltalef 10S oil or water. Epidermal cells in the A4-A7 abdominal segments of the larvae were imaged live through the cuticle using a Leica SP5 inverted confocal microscope with a 63x/1.4 oil immersion objective. Tagged fluorescent proteins were excited sequentially with 488nm and 561nm laser beams and detected with 510-540nm and 580-630nm emission filters, using Leica HyD hybrid detectors.

### Quantification of Ds Amounts at Cellular Interfaces

Ds::EGFP membrane distribution was analysed in the apical plane of ventral epidermal cells of early second stage larvae. Two juxtaposed areas of the segment (the denticulate and undenticulate regions) were imaged separately to grant sufficient resolution and subsequently merged, and maximum intensity projections of typically 4μm stacks were used to compensate for ruggedness in the denticulate region. Between 3 and 12 images from different larvae were acquired and aligned to the mediolateral axis using rows of tendon cells as reference. Ten straight lines parallel to the anteroposterior axis and 4μm wide were drawn over the images at random heights, and the profile of average fluorescence intensity along each line was plotted. Each profile displayed peaks where the line intersected cell boundaries: the fluorescence maxima were quantified using the BAR collection of ImageJ routines [**25**] and manually assigned to the respective cellular interfaces. Due to cell morphology and image noise not every line could provide a measure for each interface, therefore for every image a value of mean intensity was calculated only for cell boundaries intersected by at least 3 lines. The mean of means of all boundaries in an image was used as reference to normalise the fluorescence intensity maxima.

### Mapping of D polarity

D polarity at the plasma membrane was assessed over the whole segment by analysing a total of 594 cells from small clones expressing *d::EGFP* in the ventral epidermis of 44 different larvae. Each cell was assigned a row number and polarity: rows of cells were identifiable by proximity to conspicuous landmarks like denticles, sensory cells, and tendons with unique shape, while polarity was scored by eye based on whether D::EGFP fluorescence was exclusively on the anterior (Anterior membrane) or posterior (Posterior membrane) side of their plasma membrane, unpolarised but clearly enriched at the membrane (Uniform membrane), or homogeneously distributed in the cytoplasm (Uniform cytoplasm).

### Analysis of Microtubule Growth Direction

Orientation of growing microtubules was analysed following EB1::EGFP comets in ventral larval epidermal cells. Clonal expression of *EB1::EGFP* was necessary to avoid interference from the strong signal of underlying muscle cells, and undenticulate regions were preferred because denticles obscured the fluorescent signal. Early second stage larvae were mounted in a small drop of water ensuring their posterior spiracles were out of the liquid, and movies of individual cells were recorded at 5.16 s intervals for typically 5 min, imaging a single 0.773μm apical confocal plane. Movie frames were registered using the ImageJ plugin Stackreg [**26**] to account for slight movements of the larvae. Cells were then aligned to the mediolateral axis using the T3 row of tendon cells and rows of denticles as references, and cells situated in the right hemisegments were flipped to match the mediolateral orientation of the left hemisegment cells. Two cells, one in the A compartment (row 7 or 8) and one in the P (row −2 or −1), were selected from each of 10 larvae and pooled for blind analysis. Comets were traced manually using the ImageJ plugin MtrackJ [**27**], sampling all the visible comets within each cell for as many time points as were necessary to count 150-200 comets per cell, and angles of the comets’ trajectories relative to the anteroposterior axis of the larva were derived from the first and last time point of their tracks.

### Data Analysis

Data analysis was carried out in R 3.5.3 [**28**], using the *CircMLE* [**29**], *circular* [**30**], *DescTools* [**31**], *dplyr* [**32**], *ggplot2* [**33**], and *mosaic* [**34**] packages.

### Data Availability

Data used in **figures 4, 8, S1, S4-6** can be obtained from the University of Cambridge Open Access repository (https://doi.org/10.17863/CAM.53667)

## RESULTS

### Comparing wildtype and polarity modified larvae

#### (i) Background

In this section we reexamine and test the model as exemplified by those single cells described as “atypical” in which one face of the cell’s membrane abuts two different neighbours [**17**]. Some of these cells are multipolar and these exemplify very strongly the argument that PCP stems from a comparison between the facing membranes of a single cell. These atypical and multipolar cells are now studied in “polarity modified” larvae, in which the overall segmental polarity has been considerably modified by experiment. Unlike previously, we study the predenticles, that is denticles observed prior to the deposition of cuticle.

We compare the cell polarity of wildtype [**15, 17**] and polarity modified larvae (**figure 2**). To make the polarity modified larvae, we engineer increased expression of an active form of *ds* in T1 and T2 cells (*sr*.*Gal4 UAS*.*ectoDs* [**15**]); this changes the landscape of Ds activity, making peaks (instead of troughs, as in the wildtype) in T1 and T2. Consequently, the polarities of rows of cells 1, 2, 4 and 5, that abut T1 and T2, now point inwards; that is reversed from the wildtype (**figure 2**). The other rows, 0, 3 and 6 could also be affected because polarity can be propagated beyond the neighbouring cells [**8, 9, 15**]. To explain further how the Ds/Ft machine propagates polarity changes from cell to cell: an increase in Ds activity in cell ***a*** attracts more Ft on the facing membrane of cell ***b***. On that facing membrane more Ft tends to exclude Ds activity, enabling more Ds to accumulate on the far side of cell ***b*** which will, in turn, draw more Ft to the facing membrane of cell ***c*** [**5, 7**].

#### (ii) Atypical cells

In all larvae, the numbered cell rows are often irregular and some atypical cells may individually abut on the same side two neighbours, each with a different level of Ds activity. We compare the predenticles of atypical cells in wildtype and polarity modified larvae. In the wildtype, one posterior part of cell ***a*** in row 4 may contact a T2 neighbour with a lower Ds activity than row 3 (the associated predenticles in this region of cell ***a*** point anteriorly) and a separate part of cell ***a*** may contact a row 4 neighbour with a higher Ds activity than row 3 [**17**]. However, in the polarity modified larvae, the predenticles of nearly all cells of row 4 (typical and atypical cells) point posteriorly —this is as expected from the model because ***both*** types of posterior neighbour that can abut a row 4 cell (T2 and another row 4 cell) now have higher levels of Ds activity than the anterior neighbour, a row 3 cell (**figure 2A-C** and **table 1**). However for these polarity modified larvae, some single atypical cells of row 2 have two anterior neighbours —cells of T1 and row 2— that are higher and lower in Ds activity than the posterior neighbour of the atypical cell, respectively. Consequently, the model predicts that their associated predenticles should point forwards in that part of the cell that abuts T1 and backwards in that part of the same cell abutting row 2, and they do (**figure 2D-F** and **table 1**). There are some quantitative differences between the current data and the wildtypes we scored earlier ([**17**], see legend to **table 1)**. Nevertheless, these results, especially on the polarity modified larvae, confirm and strengthen a model of PCP in which cells in a tissue are polarised due to an underlying gradient of Ds activity.

### Direct assessment of Ds distribution in both wildtype and polarity modified larvae

We measure the native Ds distribution using a tagged Ds molecule expressed as in the wildtype. Ds accumulates as puncta in the membrane (**figure 3**, [**14, 35**]) and, presumably, the puncta contain or consist of Ds-Ft heterodimers [**36**].

We previously inferred but did not show directly a supracellular gradient in Ds activity that rises within the A compartment reaching a peak near the rear of that compartment and then falling into the P [**16**]. We therefore quantified and compared the amount of Ds localised at cell junctions in all rows of the segment in the larval ventral epidermis. These measurements do not evidence an overall gradient. However, both junctions 9/T3 and T3/10 show a higher amount of tagged Ds than the other boundaries; these junctions are located near the rear of the A compartment (**figure 4**). We applied the same quantitation technique to polarity modified larvae and found that the distribution of Ds is altered from the wildtype as expected (**figure S1**), in a way that validates our quantification technique and consequently the existence of a peak of Ds levels near the rear of the A compartment in the wildtype (**figure 4**).

### The location of Dachs

The myosin-related molecule D is a marker of polarity and localised by the Ds/Ft system [**5, 14, 37-39**]. It is usually asymmetrically distributed on a polarised cell and is thought to co-localise with the face of the cell associated with the most Ds [**14, 38, 39**]. We map D to the membranes of individual cells in the larval epidermis by making small clones of cells that express tagged D; this allows the distribution of D on a particular cell to be assessed so long as the neighbour(s) does not contain any tagged D.

We examine the distribution of D in wildtype larvae in order to reveal the molecular polarity of cells that lack denticles (**figures 5, 6**). In the P compartment, all the denticulate and undenticulate cells show a consistent molecular polarity, D being localised posteriorly in the cell. Most cells of the A compartment have the opposite polarity, with D located anteriorly. In both compartments, the location of D in the denticulate cells correlates in all cases with the denticle polarity, and this includes the cells of rows 0, 1 and 4 whose denticles point forward. The tendon cells, T1, T2 and T3 can express D but it is mostly cytoplasmic in location. The cells flanking T1 and T2 (but not T3) accumulate D at the membrane abutting the tendon cells. Unlike all the other rows, cells of row 11 show some variation in the localisation of D: about 45% localise it at the posterior cell membrane, as do cells in the P compartment; in 35% it is at the membrane but not asymmetrically localised and, in the remaining cells, D is either at the anterior or found only in the cytoplasm (**figure 6**). This means that the line where polarity changes from the A-mode to the P-mode is not at the A/P border [**16**] but anterior to it; suggesting that the second cell row anterior to the A/P cellular interface (row 10) contains the peak level of Ds activity. From that row, effects on polarity spread forwards into the A compartment and backwards into row 11 and the P compartment (see model in **figure 9**).

**Figure 9.**
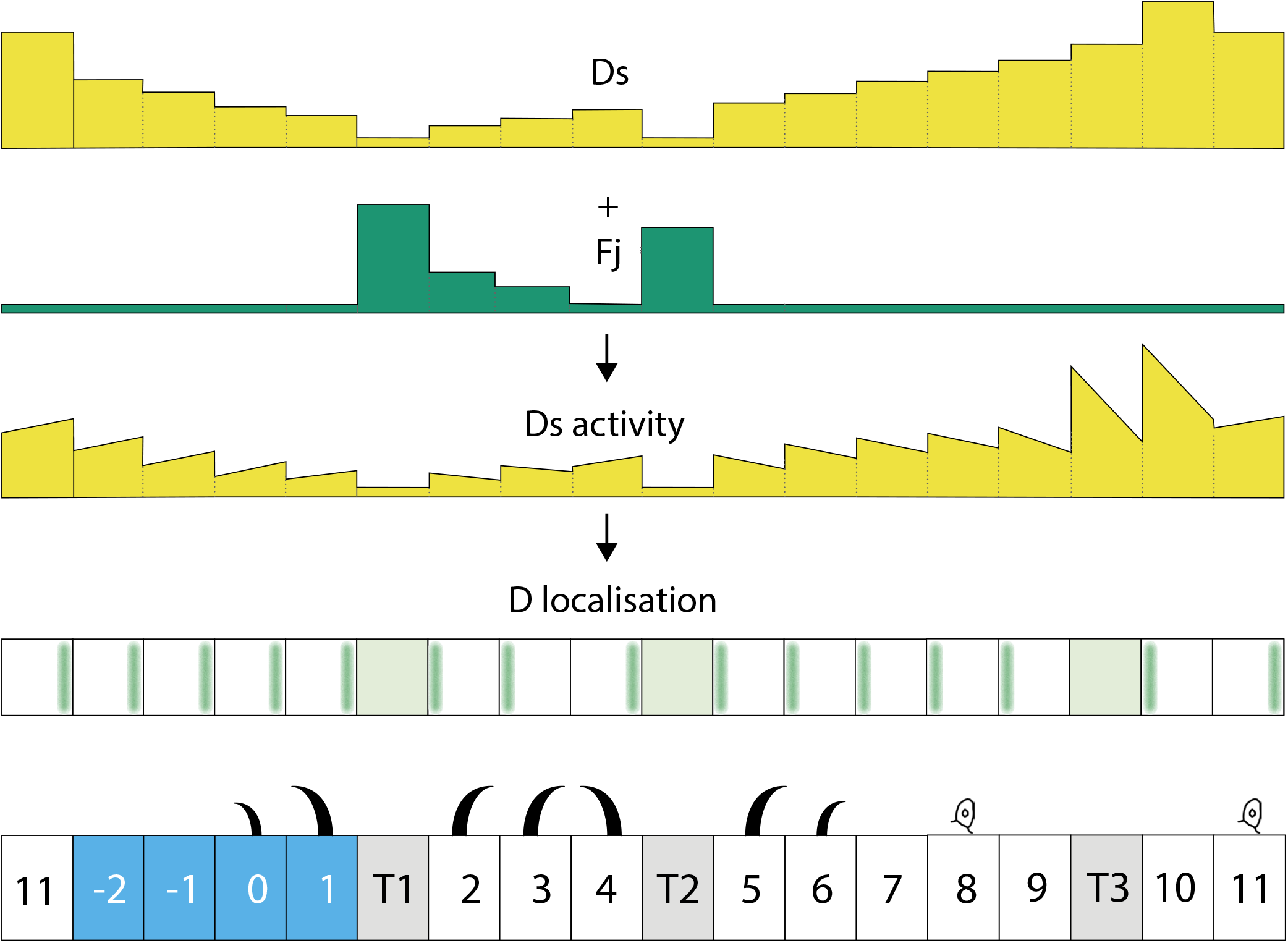
Model of Ds activity and planar cell polarity in the larval ventral epidermis. The strong Ds accumulation on both sides of T3 tendon cells (**figures 3, 4**) suggests that *ds* expression is high in T3 itself and/or its neighbours. In addition, D::EGFP clones (**figures 5, 6**) and ectopic denticles (**figure S3A**) show that polarity of row 10 points backwards, away from T3, implying that Ds activity is higher in row 11 than in T3. These two observations combined argue that *ds* expression peaks in row 10, two cells anterior to the A/P border, with Ds activity also high in T3 and row 11. Graded *ds* expression forwards and backwards from this peak together with high levels of *fj* expression in tendon cells determine the landscape of Ds activity, now extended to the undenticulate region. The Ds gradient indicated has not been confirmed, it is a speculation. Our data suggest, that if there is a pervasive gradient, it will be shallow, perhaps even more shallow than shown. The differences in Ds activity between each cell’s anterior and posterior sides orient D accumulation; D localises to the side that has the highest Ds activity and “sees” the lowest Ds activity in its neighbour. D asymmetrical distribution precisely matches the pattern of cell polarity revealed by denticles, as demonstrated by direct visualisation of tagged D in the whole segment and induction of denticles in normally naked cells. Cell 11 is shown with some ambiguity, because that is what we find (see main text). Blue shading indicates P compartment cells, grey shading tendons.

The localisation of D is not always continuous along the entire face of a cell. When the plasma membrane of one side of an atypical cell **a** abuts two separate cells, and our model implies that these two cells have different levels of Ds activity, then the D from cell **a** is localised at the interface with just one of those cells, on that part of the membrane that has most Ds activity (cells 10 and 11 in **figure 5C**, and **figure S2**, see legend). This suggests that different parts of a single cell’s membrane can compete for D.

### *ovo*-expressing clones reveal otherwise unseen polarity

Small clones that overexpress *ovo* in naked areas often produce denticles in embryos [**40, 41**]. We made marked clones in larvae and these also generally made denticles. The denticles showed a consistent orientation, pointing forwards in P and backwards in most of A, exactly mirroring the polarity pattern as identified by D localisation (**figure 7**, compare with **figure 6**). Thus, cells of row 11 at the rear of the A compartment mostly made denticles that pointed forwards (**figure 7**) as is characteristic of cells belonging to the P compartment. Just as signalled by the localisation of D, in a minority of row 11 cells, polarity was ambiguous with denticles pointing in various directions (**figure S3**). The denticles belonging to the cell row 10 anterior to row 11 always pointed backwards and denticles of the row behind row 11 (row −2 of the P compartment) always pointed forwards.

### Does the orientation of growing microtubules correlate with PCP?

We study the orientation of growing microtubules (using EB1 comets, [**42, 43**]) in the large epidermal cells of the ventral larva. Our main data is collected from identified A cells of rows 7-8 (direction of PCP is posterior) and identified P cells of rows −2 and −1 (direction of PCP is anterior; **figure 6**); the classification of the A and P cells as having opposite polarities is based on studies of the larval ventral abdomen described above. To assess the orientation of growing microtubules, we took 10 larvae, made films and studied one A and one P cell from each (**movies 1, 2**). The growing microtubules were then recorded vis-à-vis the axis of the larva by one person (SP) who was blinded to the identity of each of the 20 cells he was scoring. The orientations of about 4000 EB1 comets are shown and analysed in **figure 8**.

In the wing, the predominant alignment of the microtubules is close to the axis of PCP [**20, 44**]. By contrast, in the larval epidermal cells, in both A and P compartments, the majority of the microtubules are aligned perpendicular to the anteroposterior axis, the axis of PCP (**figure 8A,B**). To analyse our data and following the approach in the wing, the comets of the larvae are sorted into four 90 degree quadrants centred on the anteroposterior and mediolateral axes and their frequencies plotted. The quadrants are described as “anterior”, “posterior”, “medial” and “lateral” (**figure 8C,D**). The axis of PCP lies in the anteroposterior axis, but, in A compartment cells, 66% of the total angles of growth fall within the medial and lateral sectors, while in the P compartment the comparable figure is 71%. Clearly there is no overall correlation between microtubular orientation and PCP, belying the hypothesis that microtubular orientation is causal for PCP.

However, we could look for a limited correlation between the orientation of growing microtubules and the direction of PCP. For example, considering only the minority of microtubules within the anterior and posterior sectors, we find insignificant differences in polarity (**figure 8C,D**). In A cells the proportion of all microtubules that grow anteriorly is 15.8% with a 95% CI of [13.5 to 18.2] and the proportion that grow posteriorly is 18.3% [15.9 to 20.6]. In P cells it is the reverse; 16.7% grow anteriorly [14.4 to 19.1] and the proportion that grow posteriorly 12.7% [10.3 to 15.0]. There was a comparably weak bias in the medial and lateral quadrants: in A cells a larger proportion of all microtubules grow medially 34.4%[32.0 to 36.8] than laterally 31.5% [29.1 to 33.8] while the reverse bias occurs in P cells where more microtubules grow laterally 36.9% [34.5 to 39.2] than medially 33.7% [31.4 to 36.1] (**figure 8C,D**).

How uniform are the individual cells? To answer we group all the growing microtubules according to which cell (and larva) they come from and according to which of four 90 degree quadrants they fall into (**figure 8E**). Remarkably, in all sets, individual cells differ wildly from each other. Comparing the anterior versus posterior and medial versus lateral quadrants we find no strong evidence for a bias in the directions in which the microtubules grow —apart from the obvious and main finding that most of the microtubules grow more or less perpendicular to the axis of PCP.

Could there be a special subset of oriented microtubules perhaps aligned close to the anteroposterior axis, the axis of PCP, that might show a polarity bias that related to some function in planar polarity? There is no independent evidence favouring such a perspective. Nevertheless, to check we scan through the entire circumference in 22.5 degree sectors, measuring the amount of bias in the microtubules that fall within opposite pairs of sectors. There is no increase in bias in the sectors that included the axis of PCP in either the A or the P compartments, nor in nearby sectors. However, there is a local peak of bias within the A compartment: there is a significant bias in the number of growing microtubules within one pair of 22.5 degree sectors that is far away from the axis of PCP. Within the P compartment a similar peak of bias is centred near the mediolateral axis within two facing 22.5 degree sectors (**figure S4**). But note that these biases represent only 2-3% of the total population of microtubules. Thus, although we found some irregularities in the circular distribution of growing microtubules, we find no correlation with the axis of PCP.

Axelrod’s group kindly made their raw data from the pupal abdomen available to us and we treat them exactly as our larval data. Axelrod and colleagues grouped the angles of growing pupal comets into two unequal sets (two broad sectors of 170 degrees, each including the anteroposterior axis, were compared to each other, while the remaining microtubules were grouped into two narrow mediolateral sectors of 10 degrees each [**22, 23**]). But for our analysis, to conform with how data on the wing have been presented [**20, 22, 23**], and to allow a comparison with our results, we subdivided their data into four 90 degree quadrants. Even more so than in the larva, the majority of the pupal microtubules are oriented orthogonally to the axis of PCP (**figure S5A-D**): 69% of the total population of growing microtubules in the A compartment are aligned within the quadrants centred on the mediolateral axis, while in the P compartment the comparable figure is 73% (**figure S5C,D**). This finding does not fit comfortably with a hypothesis that microtubular orientation drives PCP.

Further comparison of the Axelrod group’s data on the pupa with ours on the larva show some quantitative differences. Unlike ours on the larva, their pupal data show statistically significant biases in the orientation of comets (**figure S5C,D**). In A cells the proportion of all microtubules that grow anteriorly is 12.7% with a 95% CI of [11.3 to 14.1], significantly smaller than the proportion that grow posteriorly 18.1% [16.6 to 19.5]. In P cells we see a reverse bias: 15.8% [13.3 to 18.2] grow anteriorly and 11.5% [9.1 to 13.9] posteriorly. Notably, there is a comparable and also significant bias in the medial and lateral quadrants but in the same direction in both compartments. In A cells a larger proportion of all microtubules grow laterally 38.1% [36.7 to 39.6] than medially 31.1% [29.7-32.5] and a similar bias occurs in P cells where 39.8% [37.4-42.3] grow laterally and 32.9% [30.5-35.3] grow medially (**figure S5C,D**).

We then plotted all the growing microtubules according to which pupa they came from and according to which of four 90 degree sectors they fell into (**figure S5E**). Individual pupae differ wildly from each other. In both our results on the larva and Axelrod’s results in the pupa, there is considerable inconsistency between individuals (compare **figure 8E** with **figure S5E**). Only when all cells are taken together is there any overall and significant polarity bias in Axelrod’s data.

We classified the growing microtubules in Axelrod’s data into 22.5 degree sectors and looked for an orientation bias within opposite pairs of sectors. We find examples of significant bias shown by the microtubules in various sector pairs and these are mostly not near the axis of PCP. In A cells there is a statistically significant and local peak of bias ca 60-80 degrees divergent from the axis of PCP. In P cells there is a statistically significant and local peak of bias ca 35-55 degrees divergent from the axis of PCP (**figure S4**). These observations do not fit with the conjecture that a special set of oriented microtubules, in or close to the PCP axis, might be driving planar polarity.

Dividing the data into sectors gives the impression of biases in the anteroposterior as well as in the mediolateral axes (although these are non significant in the case of the larva). But, because we suspect that subdividing the angles into sectors may lead to erroneous conclusions we investigated the distributions of the angles as a whole. We took the angular data of the A and P cells of the larva and pupal abdomen and using a maximum likelihood model approach [**29**], we found that the best fit in all four cases is to a distribution with two peaks each roughly 90 degrees divergent from the axis of PCP (**figure S6**). Unexpectedly, there are slight deviations of these peaks in the bimodal distributions; in all four distributions one of the peaks deviates 10 degrees from the mediolateral axis. Interestingly, the direction of deviation is opposite in the A cells to that in the P cells; in both sets of A cells one of the peaks is tilted 10 degree towards the posterior hemi-circumference, whereas in both sets of P cells one of the peaks is tilted 10 degrees towards the anterior hemicircumference (**figure S6**, see legend). These opposite deviations in A and P cells may be the basis of the apparent but weak biases we observe when dividing the data into four quadrants.

## DISCUSSION

### A gradient model?

In trying to understand planar cell polarity, *Drosophila* has proved the most amenable and useful experimental system. Using the *Drosophila* larva, we have built a model of how the Ds/Ft system determines the pattern of polarity in the abdominal segment [**16, 17**]. In this model the Ds/Ft system converts graded slopes in the expression levels of *ds* and *fj* into local intercellular differences in the levels of Ds activity, and into PCP without any intervention by the Stan/Fz system [**5**].

Here we have reexamined the model and extended it to those uncharted parts of the larval segment that lack denticles (**figure 9**). All the observations we have made give results that are consistent with and support the model. However it is not clear whether the model requires interactions between Ds, Ft and Fj to produce a multicellular gradient of Ds levels at the cell membranes, and expectations on this differ [**36**]. We originally proposed that the levels of Ds activity would be graded in opposite ways in the A and the P compartment and ultimately these gradients would be read out as PCP in each of the cells [**16**]. We imagined that multicellular gradients of Ds activity would persist and span the whole field of cells and this has been assumed by most [**5, 7, 45, 46**] and actually detected, locally, in the migrating larval epidermal cells in the pupa [**47**]. Alternatively, once the arrow of polarity has been established in each cell, a feedback mechanism could result in a redistribution of bridges so that, ultimately, each cell would contain the same numbers of bridges, similarly disposed— there would be no persistent multicellular gradient in Ds activity (eg [**36**]). However there would still be differences in the dispositions and orientations of Ds-Ft bridges between the opposite membranes of each cell. Our current measurements of Ds levels do not settle the matter: we did not detect a pervasive gradient of Ds, but amounts were not flat either. We found a peak in Ds level located near the rear of the A compartment near where a Ds activity gradient was predicted to summit. However a shallow Ds gradient could still exist — it might be missed because we quantify only the total Ds present in abutting pairs of membranes. This shortcoming means that the results can neither tell us the cellular provenance of the Ds we measure, nor reveal how much of it is in Ds-Ft or in Ft-Ds bridges within the apposed membranes. Thus, if any cell has a higher level of Ds, this Ds will bind more Ft in the abutting cell membrane, and, we believe, tend to exclude Ds from that abutting membrane. These effects will tend to even out the amounts of Ds in joint membranes and therefore tend to disguise any gradients, local peaks or troughs.

Could one build the segmental pattern of polarity using only a peak plus propagation, thereby managing without any initial gradient of *ds* expression? If so, a localised peak in amount of Ds at the rear of the A compartment (with a maximum in row 10) could affect polarity forwards into row 9 and beyond, and propagate backwards through row 11 into the P compartment. The single cell troughs in Ds activity in T1 and T2 would orient the polarity of the flanking cells to point away from these tendon cells. All these polarity effects would reinforce each other to make a more robust pattern. However, if there were no initial gradient of *ds* expression, only row 3 would present a problem; in order to explain why it points backwards, the trough of T1 in Ds activity would need to be deeper than that of T2 (see figure 4 in [**15**]). Perhaps it will prove important to note that the gradient model and the alternative localised peak and troughs model just outlined are not mutually exclusive and each can contain aspects of the truth.

Originally predicted to be at the A/P compartment border [**16**] we conclude now that a Ds peak occurs two cells anterior to that border, in row 10 (**figure 9**, a similar peak two cells from the A/P border has been described in the dorsal abdomen of the pupa [**47**]). This observation is supported by both D localisation and the orientation of ectopic denticles formed by *ovo*-expressing clones. There are interesting implications: the peak in Ds protein at the cell junctions is in a cell that is flanked on both sides by A compartment cells, the most posterior of which (row 11) has “P type” polarity. Why is this summit out of register with the lineage compartments? It could be that this peak is specified by a signal emanating from one compartment and crossing over to affect the next compartment. There are precedents for this kind of transgression [**48-52**]. Also, in the abdomen of the developing adult fly, Hedgehog signal spreads from the P compartment across into the A compartment and induces different types of cuticle at different distances [**53**].

Our results can best be interpreted, as others have done [**14, 37, 54**], that D acts as an eloquent marker of a cell’s polarity, is localised on the membrane with the most Ds, and acts immediately downstream of the Ds/Ft system.

### Microtubules and PCP

We have suggested [**17**] that intracellular conduits might be involved in a local comparison between facing membranes of a cell and shown here that this perspective successfully predicts which cells should become bipolar even in polarity modified larvae. But there is still no direct evidence for the conduits, and no knowledge of, if they do exist, what they are. One could imagine a set of microtubules, initiated on the membrane, that could align more or less with the anteroposterior axis and traverse the cell to meet the membrane opposite. Indeed, Uemura’s group have proposed that microtubules, oriented by the Ds/Ft system, translocate vesicles carrying PCP components such as Frizzled (Fz) and Dishevelled (Dsh) to one side of a cell to polarise it. Their hypothesis began with observations on microtubule-dependent transport of tagged proteins *in vivo* in cells of the wing disc [**19**] and was extended by the use of EB1 comets to plot microtubule polarity in the pupal wing [**20-23**]. Harumoto and colleagues studied the proximal part of the wing where they found a transient correlation, with a small majority of the microtubules growing distally, but there was no such correlation in the distal wing. Also, in *ds*^*–*^ wings, distal regions show consistently polarised microtubules (a small majority now grow proximally), although the hairs in that region still point distally [**20**]. Likewise, while some studies of the adult abdomen demonstrate a local correlation between cell polarity and the orientation of limited subsets of microtubules, PCP in other parts did not show this correlation and the authors concluded that, in those parts, polarity is determined independently of the microtubules [**23**]. We have tested the hypothesis that microtubular orientation drives PCP in the larval abdomen of *Drosophila* and there it also meets serious difficulties. The greatest of these is that most of the microtubules are aligned orthogonally to the axis of PCP (this fact is also extractable from the pupal data kindly provided by Axelrod’s group). Of the roughly 30% of all microtubules that fall into the two quadrants centred on the axis of PCP, there is a small net excess, corresponding to about 5% of the total, that could perhaps result in a net transport of vesicles in the direction of PCP. But even if this were so, more than 80% of the vesicles carrying cargo should arrive in the wrong part of the cell membrane.

Why are there apparent biases in microtubule orientation in the data? An analysis of the circular distribution of comets showed, in all the sets of data (ours and those of Axelrod’s group), a deviation of 10 degrees in one of the peaks of the bimodal distribution of the angles (**figure S6**). This deviation, plus the precise orientation of the 90 degree quadrants, may explain the apparent bias of microtubular orientation seen clearly in the Axelrod data and hinted at much more weakly in our data. How? Imagine a circular bimodal distribution composed of two separate unimodal distributions: the tails of both probability distributions would be closer and overlap more if the distance between the mean angles were reduced. In our cases, one of the tails of the distributions whose mean angles deviate by 10 degrees will decrease slightly the frequency of angles within one of the anteroposterior quadrants and concomitantly the other tail increase the frequency in the opposite anteroposterior quadrant. This deviation may have its origin in a correlation between cell shape and microtubular orientation [**44, 55, 56**] and in different cell shapes in the A and P cells; these are more obvious at or close to the A/P border [**57**].

The hypothesis of Uemura’s group which proposes that microtubules transport Fz to one side of the cell to polarise it meets an additional problem in the larval abdomen. The normal orientations of the denticles in the larva does not require input from the Stan/Fz system; indeed the Ds/Ft system appears to act alone [**7-9**]. But could oriented microtubules be involved in PCP, even without any role of the Stan/Fz system? Our results from the larval abdomen say no. We cannot exclude the possibility of a small subset of stable microtubules (undetectable because they would not bind EB1), aligned with the anteroposterior axis and strongly biased in polarity, in the pupal or larval abdomens (or proximodistal axis in the wing). There is no evidence for such microtubules, but if they exist their number and bias in orientation must be strong enough to overcome the moving of vesicles on the unbiased dynamic microtubules we have studied.

### Conclusions

We have enhanced our present model of how the Ds/Ft system generates the intricate polarity of the larval segment. The key element of this model is that each cell compares its neighbours and is polarised (and points its denticles) towards the cell presenting the most Ds activity. This hypothesis gains more support from our new results on the multipolarity of single cells. But we have not found out how the comparison is made: an attractive hypothesis by others was that oriented microtubules are the critical agent, but, if we interrogate our data for biases in polarity within all the growing microtubules, or if we select subsets of microtubules whose orientations are related to the axis of PCP, we do not find evidence for a link between microtubular polarity and the polarity of the denticles (the “direction” of PCP). Using two different methods we demonstrated that undeticulated cells are also polarised and their polarity is as the model predicts, and that the point where the amount of Ds is, presumably, highest and from where, like a watershed divide, polarity diverges, is two cells away from the compartment border. We looked to demonstrate the predicted multicellular gradient of Ds but, possibly because of an insufficiency in our methods, we only found a localised peak (at the rear of the A compartment as the model requires). Thus, if there is a multicellular gradient of Ds activity, it must be very shallow. There’s still much to do; still so much to learn.

## ACKNOWLEDGEMENTS

We thank Jeffrey Axelrod and Katherine Sharp for kindly sharing data from the Axelrod group (published in [**22, 23**]), and David Strutt, Eduardo Moreno, and the Bloomington Stock Center for flies.

## COMPETING INTERESTS

The authors declare that no competing interests exist.

## FUNDING

Our work was supported by Wellcome Investigator Award 107060 to PAL.

---

**Movie 1**. Film of microtubule dynamics in a representative larval A cell. EB1::GFP comets in a row 7 cell from the right hemisegment imaged for 4 minutes at 5.16s intervals. Juxtaposed movie shows manual tracing of 200 comet trajectories over the entire surface of the cell. Anterior is to the left, medial is down. Scale bar: 5**μ**m.

**Movie 2**. Film of microtubule dynamics in a representative larval P cell. EB1::GFP comets in a row −1 cell from the left hemisegment imaged for 4 minutes at 5.16 s intervals. Juxtaposed movie shows manual tracing of 200 comet trajectories over the entire surface of the cell. Anterior is to the left, medial is up. Scale bar: 5**μ**m.

## SUPPLEMENTARY FIGURE LEGENDS

**Figure S1.**
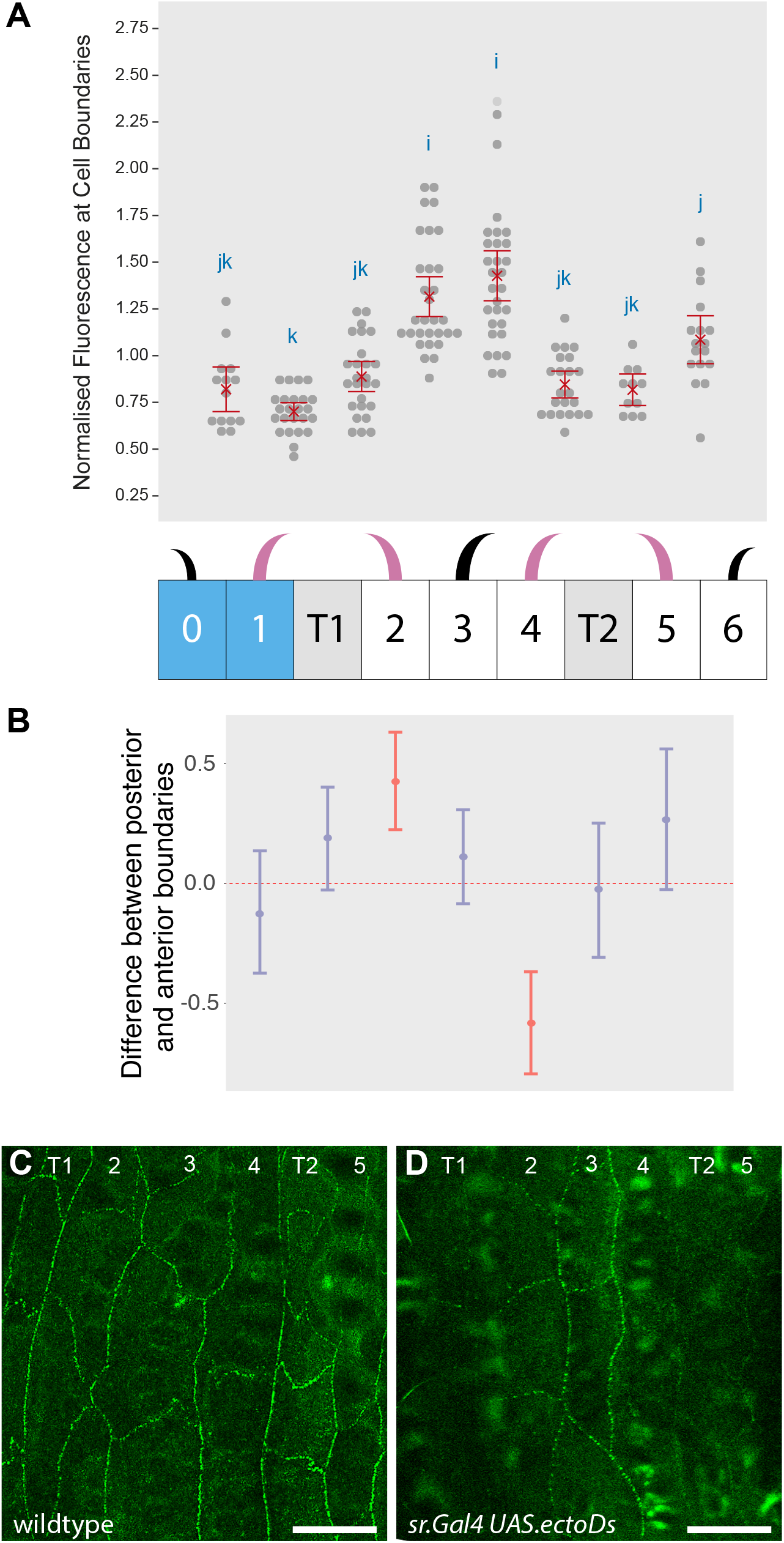
Quantitation of Ds levels at cellular interfaces in polarity modified larvae. (**A**) Dot plot, diagram of denticle polarity, and (**B**) pairwise comparisons are presented as in **figure 4**. Data are pooled from 3 images of larvae where overexpression of untagged Ds is specifically driven in tendons and changes the polarity of adjacent denticle cells (see **figure 2**). Ds distribution in (**C**) wild type (a detail from **figure 3A**) and (**D**) polarity modified larvae (*sr*.*Gal4 UAS*.*ectoDs*) is clearly different, reflecting the predicted changes in the landscape of Ds activity. For example in (**D**), more untagged Ds in T1 attracts more Ft molecules in row 2 cells to the T1/2 boundary, consequently displacing the row 2 endogenous, tagged Ds to the 2/3 boundary and raising fluorescence on that interface. The same effect emanating anteriorly from T2 raises Ds fluorescence at the 3/4 boundary. As expected, Ds amounts on the 2/3 and 3/4 boundaries are significantly higher than on the surrounding boundaries, arguing that the method is capable of detecting cellular interfaces with raised Ds activity. Attempts to relate observed polarity of a cell with the localisation of Ds at its membranes are compromised because we cannot determine how much Ds each of the two abutting cells is contributing to their joint membrane. It is interesting to note that overexpressing ectoDs in the tendon cells has no significant effect on the amount of tagged Ds in 1/T1 or T2/5 boundaries. We think that is due to the several cells anterior to row 1 and the several cells posterior to row 6 dampening the effects. Scale bars: 20μm.

**Figure S2.**
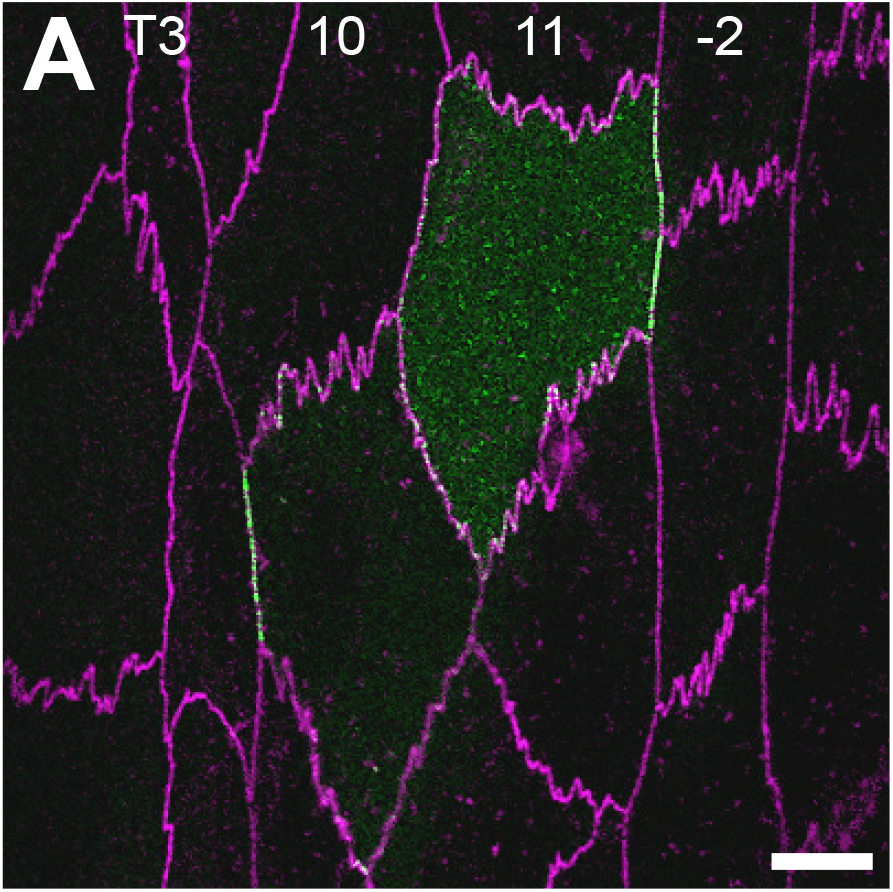
D localisation on limited parts of the plasma membrane. (**A**) Row 10 and 11 cells from a wildtype larva expressing *d::EGFP*, with cell outlines marked in magenta by DE-cad::tomato (see **figure 5C** for single EGFP channel). D is on just one side of each cell, but its localisation at the plasma membrane is not continuous: the row 10 cell accumulates D on the anterior membrane only where it confronts a T3 cell, not where it faces other row 10 cells; the row 11 cell has D localised at its posterior face, but only where it contacts row −2 cells. Scale bar: 10μm.

**Figure S3.**
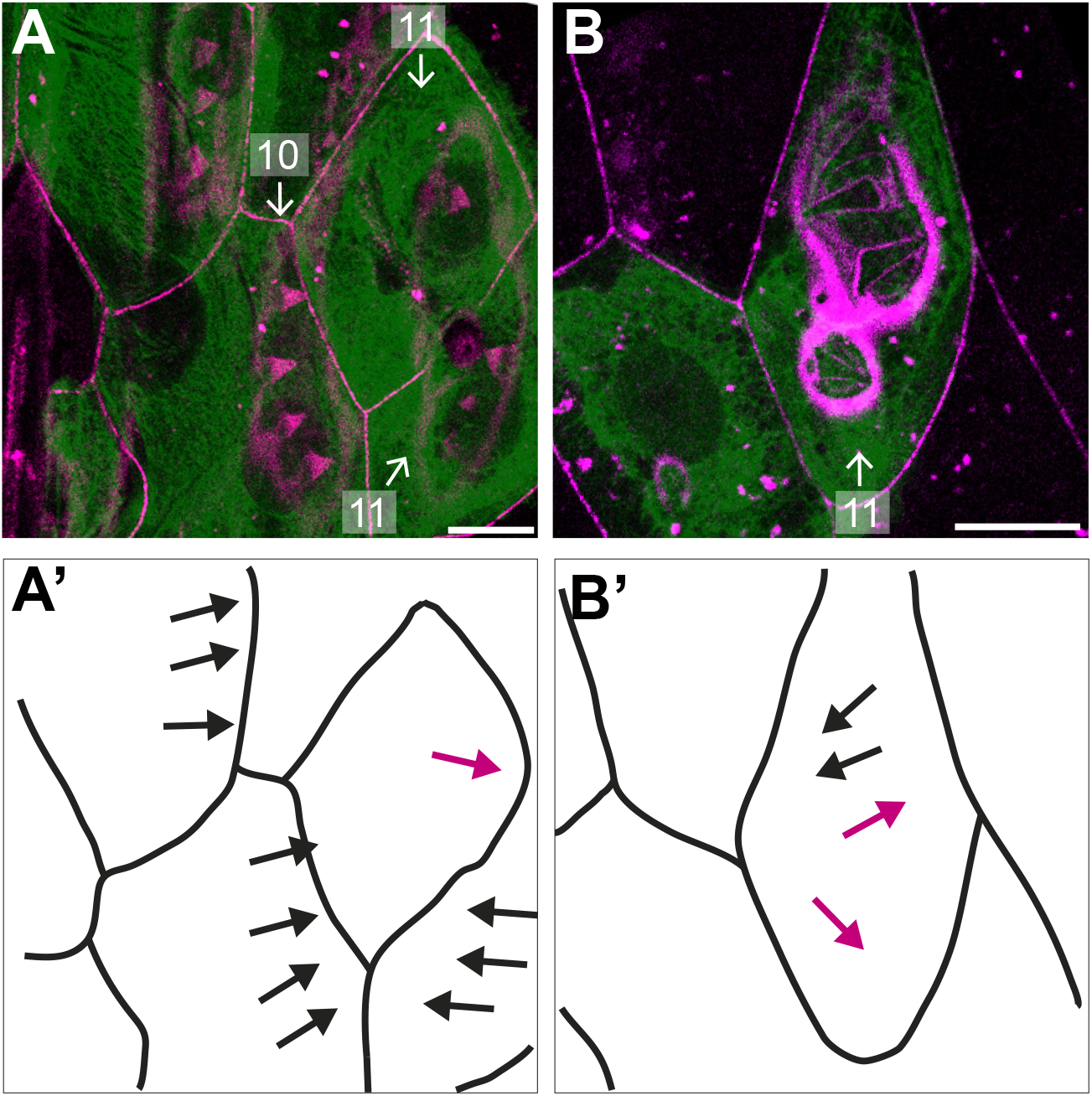
Unusual *ovo*-expressing clones with ambiguous polarity in row 11 cells. (**A,B**) Clones marked with EGFP and producing ectopic denticles in rows 10 and 11. *DE-cad::tomato* (magenta) labels cell boundaries and denticles, which in this area can be tenuous and hard to discern. (**A’,B’**) Schemes of cell outlines and denticle orientation; denticles with uncharacteristic polarity are highlighted in red. (**A,A’**) Denticles pointing in opposite directions in two contiguous row 11 cells; all denticles in the neighbouring row 10 cells, point backwards. (**B,B’**) Denticles pointing in mixed directions within a single row 11 cell. Scale bars: 10μm.

**Figure S4.**
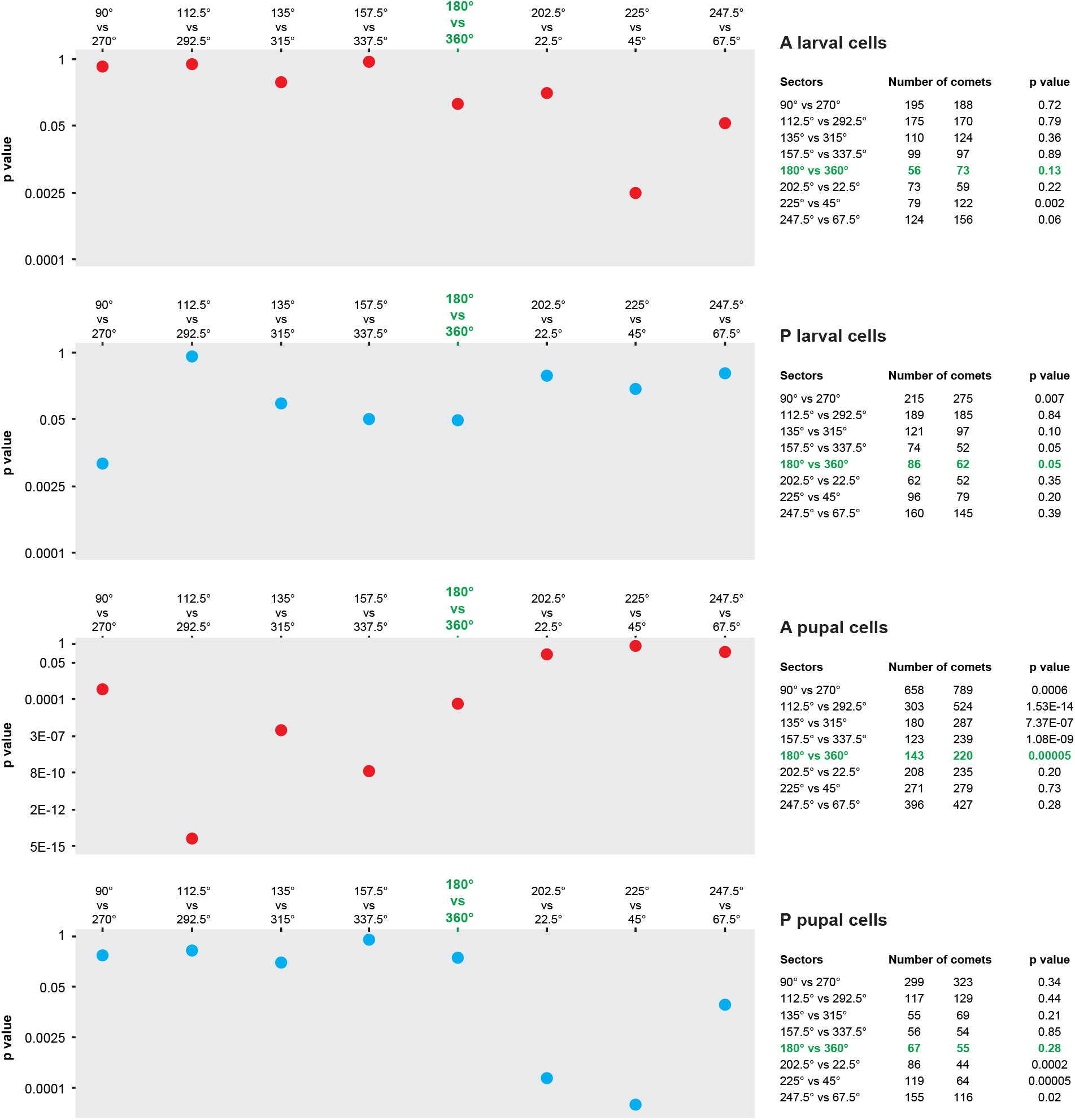
Local polarity biases in microtubule growth. P values of chi-squared tests between numbers of comets whose orientation falls in opposite 22.5 degree sectors. Tables display the number of comets per sector and p values for larval and pupal sets of A and P cells. Sectors centred on the anteroposterior axis are highlighted in green.

**Figure S5.**
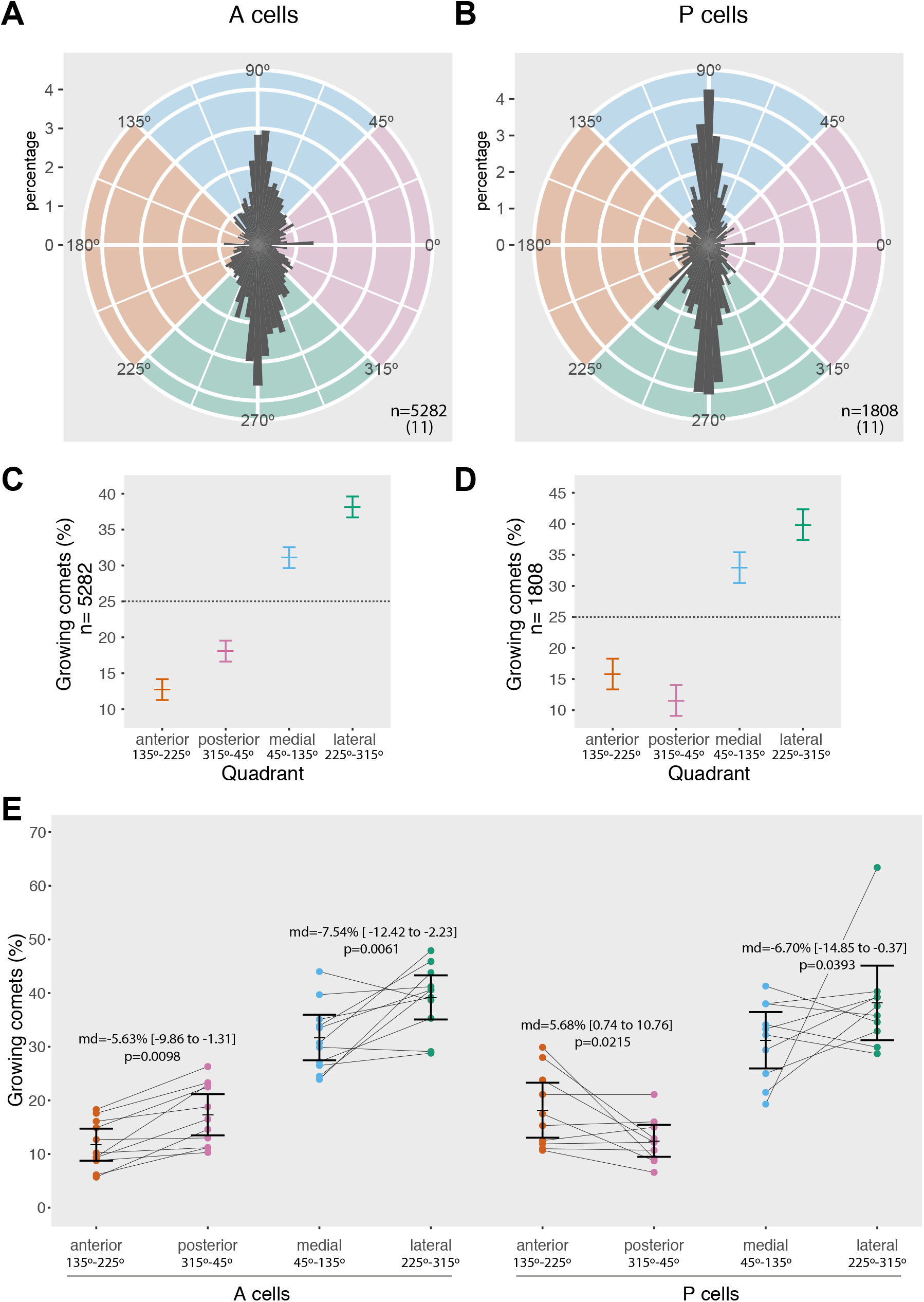
Analysis of microtubule polarity in cells of the pupal abdomen, based on raw data kindly provided by the Axelrod group. (**A-E**) Rose diagrams of microtubule growth distribution, frequencies of comet orientation, and dot plot of microtubule direction in individual cells are presented as in **figure 8**. (**A,C,E**) Anterior pupal cells, (**B,D,E**) posterior pupal cells. n indicates total number of comets analysed, from the amount of pupae specified in parenthesis. Unlike ours, the data acquired by Axelrod’s group contain no information about which hemisegment they were sampled from; comet orientation is still classified as medial and lateral to facilitate comparison with our results, however these categories should be considered with caution. Note that, in contrast with larval data where differences between the frequencies of comets in opposite quadrants are very weak (**figure 8C,D**), in pupae there are significant biases in the proportion of anteroposteriorly and mediolaterally growing microtubules (see non-overlapping confidence intervals in **C** and **D**).

**Figure S6.**
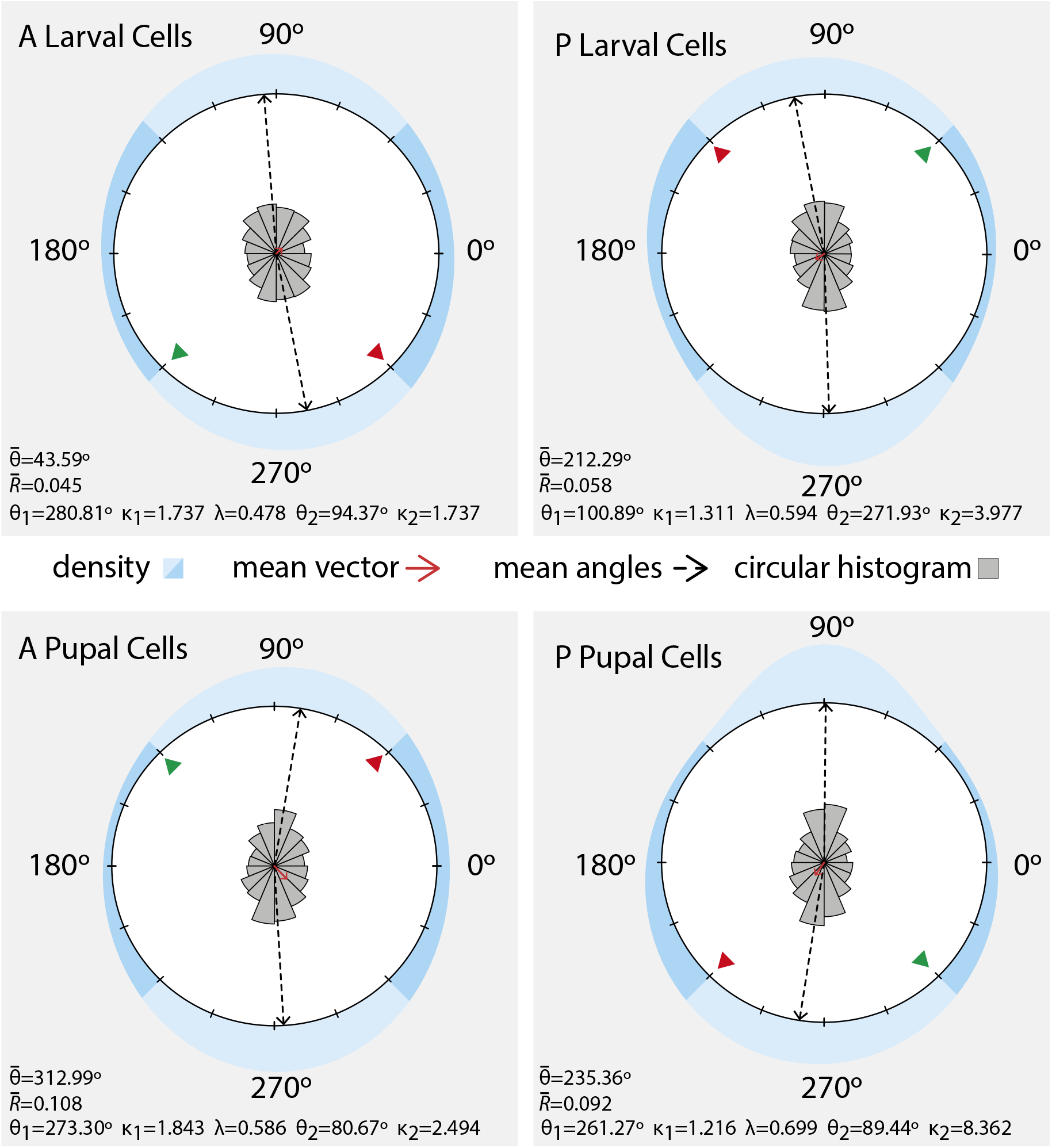
Maximum likelihood best models of microtubule angular distributions. Using a maximum likelihood approach [**34**] we plot the angular distribution of all growing microtubules and the best fit is to bimodal distributions with two peaks near 180 degrees apart in the mediolateral axis. The distribution densities are shown in blue (darker blue representing the anterior and posterior 90 degree quadrants). A circular histogram (bin size 22.5 degree) of the angle data is at the centre of each plot in grey. The mean vector is shown in red and the two mean angles are shown with discontinuous arrows. The mean values (θ), concentration parameters (κ), proportional size of the first distribution (λ), mean vector angle 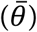 and dispersion 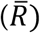 are shown below each plot. A deviation of 10 degrees in one of the peaks of the distribution from the true mediolateral axis is enough to create a difference in the density area of the anterior and posterior quadrants. In both larval and pupal sets of A cells the area of the posterior quadrant density closest to the deviated peak is slightly bigger (red arrowhead) than the anterior one (green arrowhead). In both larval and pupal sets of P cells the area of the anterior quadrant is slightly bigger (red arrowhead) than the posterior one (green arrowhead).

## Notes

### Competing Interest Statement

The authors have declared no competing interest.

## REFERENCES

1 Goodrich, L. V., Strutt, D. (2011) Principles of planar polarity in animal development. Development. 138, 1877–1892. (10.1242/dev.054080)

2 Henderson, D. J., Long, D. A., Dean, C. H. (2018) Planar cell polarity in organ formation. Curr. Opin. Cell Biol. 55, 96–103. (10.1016/j.ceb.2018.06.011)

3 Devenport, D. (2016) Tissue morphodynamics: Translating planar polarity cues into polarized cell behaviors. Semin. Cell Dev. Biol. 55, 99–110. (10.1016/j.semcdb.2016.03.012)

4 Butler, M. T., Wallingford, J. B. (2017) Planar cell polarity in development and disease. Nat. Rev. Mol. Cell Biol. 18, 375–388. (10.1038/nrm.2017.11)

5 Lawrence, P. A., Casal, J. (2018) Planar cell polarity: two genetic systems use one mechanism to read gradients. Development. 145, (10.1242/dev.168229)

6 Carvajal-Gonzalez, J. M., Mlodzik, M. (2014) Mechanisms of planar cell polarity establishment in Drosophila. F1000Prime Rep. 6, 98. (10.12703/P6-98)

7 Casal, J., Lawrence, P. A., Struhl, G. (2006) Two separate molecular systems, Dachsous/Fat and Starry night/Frizzled, act independently to confer planar cell polarity. Development. 133, 4561–4572. (10.1242/dev.02641)

8 Donoughe, S., DiNardo, S. (2011) dachsous and frizzled contribute separately to planar polarity in the Drosophila ventral epidermis. Development. 138, 2751–2759. (10.1242/dev.063024)

9 Repiso, A., Saavedra, P., Casal, J., Lawrence, P. A. (2010) Planar cell polarity: the orientation of larval denticles in Drosophila appears to depend on gradients of Dachsous and Fat. Development. 137, 3411–3415. (10.1242/dev.047126)

10 Simon, M. A., Xu, A., Ishikawa, H. O., Irvine, K. D. (2010) Modulation of Fat:dDchsous binding by the cadherin domain kinase Four-jointed. Curr. Biol. 20, 811–817. (10.1016/j.cub.2010.04.016)

11 Ishikawa, H. O., Takeuchi, H., Haltiwanger, R. S., Irvine, K. D. (2008) Four-jointed is a Golgi kinase that phosphorylates a subset of cadherin domains. Science. 321, 401–404. (10.1126/science.1158159)

12 Brittle, A., Repiso, A., Casal, J., Lawrence, P. A., Strutt, D. (2010) Four-jointed modulates growth and planar polarity by reducing the affinity of Dachsous for Fat. Curr. Biol. 20, 803–810. (10.1016/j.cub.2010.03.056)

13 Ambegaonkar, A. A., Pan, G., Mani, M., Feng, Y., Irvine, K. D. (2012) Propagation of Dachsous-Fat planar cell polarity. Curr. Biol. 22, 1302–1308. (10.1016/j.cub.2012.05.049)

14 Brittle, A., Thomas, C., Strutt, D. (2012) Planar polarity specification through asymmetric subcellular localization of Fat and Dachsous. Curr. Biol. 22, 907–914. (10.1016/j.cub.2012.03.053)

15 Saavedra, P., Brittle, A., Palacios, I. M., Strutt, D., Casal, J., Lawrence, P. A. (2016) Planar cell polarity: the Dachsous/Fat system contributes differently to the embryonic and larval stages of Drosophila. Biol. Open. 5, 397–408. (10.1242/bio.017152)

16 Casal, J., Struhl, G., Lawrence, P. A. (2002) Developmental compartments and planar polarity in Drosophila. Curr. Biol. 12, 1189–1198. (10.1016/s0960-9822(02)00974-0)

17 Rovira, M., Saavedra, P., Casal, J., Lawrence, P. A. (2015) Regions within a single epidermal cell of Drosophila can be planar polarised independently. eLife. 4, e06303. (10.7554/eLife.06303)

18 Saavedra, P., Vincent, J. P., Palacios, I. M., Lawrence, P. A., Casal, J. (2014) Plasticity of both planar cell polarity and cell identity during the development of Drosophila. eLife. 3, e01569. (10.7554/eLife.01569)

19 Shimada, Y., Yonemura, S., Ohkura, H., Strutt, D., Uemura, T. (2006) Polarized transport of Frizzled along the planar microtubule arrays in Drosophila wing epithelium. Dev. Cell. 10, 209–222. (10.1016/j.devcel.2005.11.016)

20 Harumoto, T., Ito, M., Shimada, Y., Kobayashi, T. J., Ueda, H. R., Lu, B., Uemura, T. (2010) Atypical cadherins Dachsous and Fat control dynamics of noncentrosomal microtubules in planar cell polarity. Dev. Cell. 19, 389–401. (10.1016/j.devcel.2010.08.004)

21 Matis, M., Russler-Germain, D. A., Hu, Q., Tomlin, C. J., Axelrod, J. D. (2014) Microtubules provide directional information for core PCP function. eLife. 3, e02893. (10.7554/eLife.02893)

22 Olofsson, J., Sharp, K. A., Matis, M., Cho, B., Axelrod, J. D. (2014) Prickle/spiny-legs isoforms control the polarity of the apical microtubule network in planar cell polarity. Development. 141, 2866–2874. (10.1242/dev.105932)

23 Sharp, K. A., Axelrod, J. D. (2016) Prickle isoforms control the direction of tissue polarity by microtubule independent and dependent mechanisms. Biol. Open. 5, 229–236. (10.1242/bio.016162)

24 Thurmond, J., Goodman, J., Strelets, V., Attrill, H., Gramates, L., Marygold, S., Matthews, B., Millburn, G., Antonazzo, G., Trovisco, V., et al. 2019 FlyBase 2.0: the next generation. Nucleic Acids Res. 47, D759–D765. (10.1093/nar/gky1003)

25 Ferreira, T., Hiner, M., Rueden, C., Miura, K., Eglinger, J., Chef, B. BAR 1.5.1. (2017) Available from: https://doi.org/10.5281/zenodo.495245

26 Thevenaz, P., Ruttimann, U. E., Unser, M. (1998) A pyramid approach to subpixel registration based on intensity. IEEE T. Image Process. 7, 27–41. (10.1109/83.650848)

27 Meijering, E., Dzyubachyk, O., Smal, I. (2012) Methods for cell and particle tracking. Methods Enzymol. 504, 183–200. (10.1016/b978-0-12-391857-4.00009-4)

28 R Core Team. R: A Language and Environment for Statistical Computing. Vienna, Austria: R Foundation for Statistical Computing 2019.

29 Fitak, R. R., Johnsen, S. (2017) Bringing the analysis of animal orientation data full circle: model-based approaches with maximum likelihood. J. Exp. Biol. 220, 3878–3882. (10.1242/jeb.167056)

30 Agostinelli, C., Lund. R package ‘circular’: Circular Statistics (version 0.4-93). (2017) Available from: https://r-forge.r-project.org/projects/circular/

31 Signorell, A., multitple authors. DescTools: Tools for Descriptive Statistics. (2019) Available from: https://cran.r-project.org/package=DescTools

32 Wickham, H., François, R., Henry, L., Müller, K. dplyr: A Grammar of Data Manipulation. (2019) Available from: https://CRAN.R-project.org/package=dplyr

33 Wickham, H. (2016) ggplot2: Elegant Graphics for Data Analysis. Springer-Verlag New York.

34 Pruim, R., Kaplan, D. T., Horton, N. J. (2017) The mosaic Package: Helping Students to ‘Think with Data’ Using R. R J. 9, 77–102.

35 Ma, D., Yang, C. H., McNeill, H., Simon, M. A., Axelrod, J. D. (2003) Fidelity in planar cell polarity signalling. Nature. 421, 543–547. (10.1038/nature01366)

36 Hale, R., Brittle, A. L., Fisher, K. H., Monk, N. A., Strutt, D. (2015) Cellular interpretation of the long-range gradient of Four-jointed activity in the Drosophila wing. eLife. 4, e05789. (10.7554/eLife.05789)

37 Bosveld, F., Bonnet, I., Guirao, B., Tlili, S., Wang, Z., Petitalot, A., Marchand, R., Bardet, P. L., Marcq, P., Graner, F., et al. 2012 Mechanical control of morphogenesis by Fat/Dachsous/Four-jointed planar cell polarity pathway. Science. 336, 724–727. (10.1126/science.1221071)

38 Mao, Y., Rauskolb, C., Cho, E., Hu, W. L., Hayter, H., Minihan, G., Katz, F. N., Irvine, K. D. (2006) Dachs: an unconventional myosin that functions downstream of Fat to regulate growth, affinity and gene expression in Drosophila. Development. 133, 2539–2551. (10.1242/dev.02427)

39 Rogulja, D., Rauskolb, C., Irvine, K. D. (2008) Morphogen control of wing growth through the Fat signaling pathway. Dev. Cell. 15, 309–321. (10.1016/j.devcel.2008.06.003)

40 Delon, I., Chanut-Delalande, H., Payre, F. (2003) The Ovo/Shavenbaby transcription factor specifies actin remodelling during epidermal differentiation in Drosophila. Mech. Dev. 120, 747–758. (10.1016/s0925-4773(03)00081-9)

41 Walters, J. W., Dilks, S. A., DiNardo, S. (2006) Planar polarization of the denticle field in the Drosophila embryo: roles for Myosin II (zipper) and fringe. Dev. Biol. 297, 323–339. (10.1016/j.ydbio.2006.04.454)

42 Akhmanova, A., Steinmetz, M. O. (2008) Tracking the ends: a dynamic protein network controls the fate of microtubule tips. Nat. Rev. Mol. Cell Biol. 9, 309–322. (10.1038/nrm2369)

43 Schuyler, S. C., Pellman, D. (2001) Microtubule “plus-end-tracking proteins”: The end is just the beginning. Cell. 105, 421–424. (10.1016/s0092-8674(01)00364-6)

44 Gomez, J. M., Chumakova, L., Bulgakova, N. A., Brown, N. H. (2016) Microtubule organization is determined by the shape of epithelial cells. Nat. Commun. 7, 13172. (10.1038/ncomms13172)

45 Fulford, A. D., McNeill, H. (2019) Fat/Dachsous family cadherins in cell and tissue organisation. Curr. Opin. Cell Biol. 62, 96–103. (10.1016/j.ceb.2019.10.006)

46 Matis, M., Axelrod, J. D. (2013) Regulation of PCP by the Fat signaling pathway. Genes Dev. 27, 2207–2220. (10.1101/gad.228098.113)

47 Arata, M., Sugimura, K., Uemura, T. (2017) Difference in Dachsous levels between migrating cells coordinates the direction of collective cell migration. Dev. Cell. 42, 479–497 e410. (10.1016/j.devcel.2017.08.001)

48 Basler, K., Struhl, G. (1994) Compartment boundaries and the control of Drosophila limb pattern by Hedgehog protein. Nature. 368, 208–214. (10.1038/368208a0)

49 Diaz-Benjumea, F. J., Cohen, S. M. (1995) Serrate signals through Notch to establish a Wingless-dependent organizer at the dorsal/ventral compartment boundary of the Drosophila wing. Development. 121, 4215–4225.

50 Doherty, D., Feger, G., Younger-Shepherd, S., Jan, L. Y., Jan, Y. N. (1996) Delta is a ventral to dorsal signal complementary to Serrate, another Notch ligand, in Drosophila wing formation. Genes Dev. 10, 421–434. (10.1101/gad.10.4.421)

51 Lawrence, P. A., Struhl, G. (1996) Morphogens, compartments, and pattern: lessons from Drosophila? Cell. 85, 951–961. (10.1016/s0092-8674(00)81297-0)

52 Tabata, T., Takei, Y. (2004) Morphogens, their identification and regulation. Development. 131, 703–712. (10.1242/dev.01043)

53 Struhl, G., Barbash, D. A., Lawrence, P. A. (1997) Hedgehog organises the pattern and polarity of epidermal cells in the Drosophila abdomen. Development. 124, 2143–2154.

54 Ambegaonkar, A. A., Irvine, K. D. (2015) Coordination of planar cell polarity pathways through Spiny-legs. eLife. 4, e09946. (10.7554/eLife.09946)

55 Picone, R., Ren, X., Ivanovitch, K. D., Clarke, J. D., McKendry, R. A., Baum, B. (2010) A polarised population of dynamic microtubules mediates homeostatic length control in animal cells. PLoS Biology. 8, e1000542. (10.1371/journal.pbio.1000542)

56 Singh, A., Saha, T., Begemann, I., Ricker, A., Nusse, H., Thorn-Seshold, O., Klingauf, J., Galic, M., Matis, M. (2018) Polarized microtubule dynamics directs cell mechanics and coordinates forces during epithelial morphogenesis. Nat. Cell Biol. 20, 1126–1133. (10.1038/s41556-018-0193-1)

57 Umetsu, D., Aigouy, B., Aliee, M., Sui, L., Eaton, S., Julicher, F., Dahmann, C. (2014) Local increases in mechanical tension shape compartment boundaries by biasing cell intercalations. Curr. Biol. 24, 1798–1805. (10.1016/j.cub.2014.06.052)

58 Blair, S. S. (1995) Compartments and appendage development in Drosophila. BioEssays. 17, 299–309. (10.1002/bies.950170406)

59 Sison, C. P., Glaz, J. (1995) Simultaneous confidence intervals and sample size determination for multinomial proportions. J. Am. Stat. Assoc. 90, 366–369. (10.1080/01621459.1995.10476521)

